# Insights from Incorporating Quantum Computing into Drug Design Workflows

**DOI:** 10.1101/2022.07.11.499644

**Authors:** Bayo Lau, Prashant S. Emani, Jackson Chapman, Lijing Yao, Tarsus Lam, Paul Merrill, Jonathan Warrell, Mark B. Gerstein, Hugo Y.K. Lam

## Abstract

While many quantum computing (QC) methods promise theoretical advantages over classical counterparts, quantum hardware remains limited. Exploiting near-term QC in computer-aided drug design (CADD) thus requires judicious partitioning between classical and quantum calculations. We present HypaCADD, a hybrid classical-quantum workflow for finding ligands binding to proteins, while accounting for genetic mutations. We explicitly identify modules of our drug design workflow currently amenable to replacement by QC: non-intuitively, we identify the mutation-impact predictor as the best candidate. HypaCADD thus combines classical docking and molecular dynamics with quantum machine learning (QML) to infer the impact of mutations. We present a case study with the SARS-CoV-2 protease and associated mutants. We map a classical machine-learning module onto QC, using a neural network constructed from qubit-rotation gates. We have implemented this in simulation and on two commercial quantum computers. We find that the QML models can perform on par with, if not better than, classical baselines. In summary, HypaCADD offers a successful strategy for leveraging QC for CADD.

## Introduction

Rapid advancement of biotechnologies is enabling an unprecedented rate of data generation in the biomedical domain, with game-changing applications if we can harness even some of the multi-scale, many-body biological complexity. Computational biology has provided critical contributions to the biological domain, for example, from protein folding, to genome assembly, to variant detection, etc. Core computational biology algorithms have thus far been successful in pushing the limit of classical computers, with horizontal scaling across multiple computing units limited mainly by network speed, and vertical scaling in a single computing unit limited by Moore’s Law. One important application of biotechnology is drug discovery, with costly experimental work benefiting from, for example, the lead search capabilities of Computer-Aided Drug Design (CADD). For such combinatorial, many-body problems, modern techniques are limited by computational hardware in both the information content of a model as well as the system size for a given model that can be computed with reasonable computational resources (Sliwoski et. al., 2014).

Quantum computing (QC) provides an alternative computing paradigm. Quantum algorithms promise efficient solutions to problems that are difficult through classical computing (Feynman, 1982; https://quantumalgorithmzoo.org/), and could be applicable to complicated biochemical systems. Although general-purpose, fault-tolerant, large-scale quantum computing technologies are still in the works, QC technology has already enabled promising simulation of non-trivial Hamiltonians (Smith et. al., 2019). Current noisy intermediate-scale quantum (NISQ)-era (Preskill, 2019) quantum hardware, while significantly limited by noise and short coherence time, is available even through cloud service providers (https://aws.amazon.com/braket/; https://azure.microsoft.com/en-us/services/quantum/). While practical quantum supremacy is still under debate, there is certainly enough evidence that quantum computing could be better than analogous classical computing in some aspects in the near future. In fact, many institutions have started to explore QC as applied to drug discovery, with near-term focuses on selected tasks such as lead optimization or compound screening (Zinner et. al., 2021).

Many of the advantages of QC are related to the fact that quantum bits, or qubits, provide a state space that is exponentially larger than that of classical counterparts. Thus, while a classical bit has two discrete states, 0 and 1, a qubit can be treated as a rotatable vector that can be represented by two continuously varying angles. Moreover, by interfering with each other like waves, these qubits can take on joint configurations inaccessible to classical systems. These behaviors may lend QC advantages over classical computing in terms of speed, and potentially even representation of data (Emani et. al., 2021). However, currently available commercial quantum computers, based on the number of available qubits, limit the size of problems that can be mapped and solved. This is especially true for the logic-gate systems, which perform logical operations via sequential qubit-rotation gates. Available numbers of qubits are on the order of ∼100 (Ball, 2021; https://azure.microsoft.com/en-us/services/quantum/; https://www.ibm.com/quantum-computing), at the higher end. Furthermore, many published QC algorithms were designed based on a putative quantum equivalent of RAM, or qRAM (Giovannetti et. al., 2008), where superpositions of qubits can be directly queried. Current lack of commercially available realizations of qRAM implies near-term use of QC algorithms has to rely on classical memory. As such, any real-world applications of QC would have to judiciously divide tasks between classical and QC capabilities. These “hybrid” approaches would have to first be evaluated for feasibility (i.e. solving the problem to reasonable accuracy), and subsequently for potential quantum advantages over purely classical equivalents. The question of feasibility is especially important in biological problems where the structure of the data is complex, and has recently been considered for the case of mRNA codon optimization (Fox et. al., 2021).

Here, we explore the incorporation of available QC into an otherwise classical computational screening workflow. We take this hybrid approach with two primary aims: (a) demonstrating how quantum computing can be combined with classical computing to create a tool that solves real-world, multi-dimensional biomedical problems; and (b) comparing multiple quantum machine learning algorithms to published classical counterparts, both in simulation and on actual quantum computers.

Specifically, we demonstrate a hybrid computing approach, named HypaCADD, applied towards CADD. HypaCADD is a hyperscale computational pipeline that integrates large-scale genomics and protein structure data for drug discovery. It consists of highly computationally intensive applications, such as molecular docking, binding affinity prediction, molecular dynamics, and machine-learning-based lead search and optimization. To prove the readiness and potential utility of quantum computing in drug discovery research, the pipeline uses quantum machine learning (QML) for predicting mutational effects on drug binding. It compares the quantum and classical computing results, and demonstrates the consistency between running in simulation and on real quantum computers by Rigetti (https://www.rigetti.com) and IBM (https://www.ibm.com/quantum-computing). We choose a straightforward form of hybridity: the data is preprocessed on a classical system and fed to quantum simulators and/or hardware. We identify an independent module of the workflow (Figure 1) to convert to a quantum analogue, namely the module associated with calculating the impact of amino acid mutations on ligand binding affinity. The advantages of this approach are twofold. First, we utilize the published results of a classical machine learning approach (Wang et. al. 2019) for careful validation of the quantum methods. Second, we completely replace this module in a tractable manner for current quantum simulators and hardware, thereby applying state-of-the-art QC to an important component of the drug-design pipeline. Downstream, we aim to start replacing, partially or even fully, some of the more computationally complex components, such as the molecular docking and molecular dynamics algorithms. As QC continues to advance, it can also be used for virtual screening with larger-size molecules or for more accurate and faster molecular dynamics simulation. For now, we find that many of the quantum algorithms we use for the mutation-impact module perform on par with their classical counterparts. HypaCADD thus affirms the value of leveraging a hybrid approach to make quantum computing readily accessible and meaningful to biomedical scientists in solving challenging drug discovery problems.

**Figure 1.**
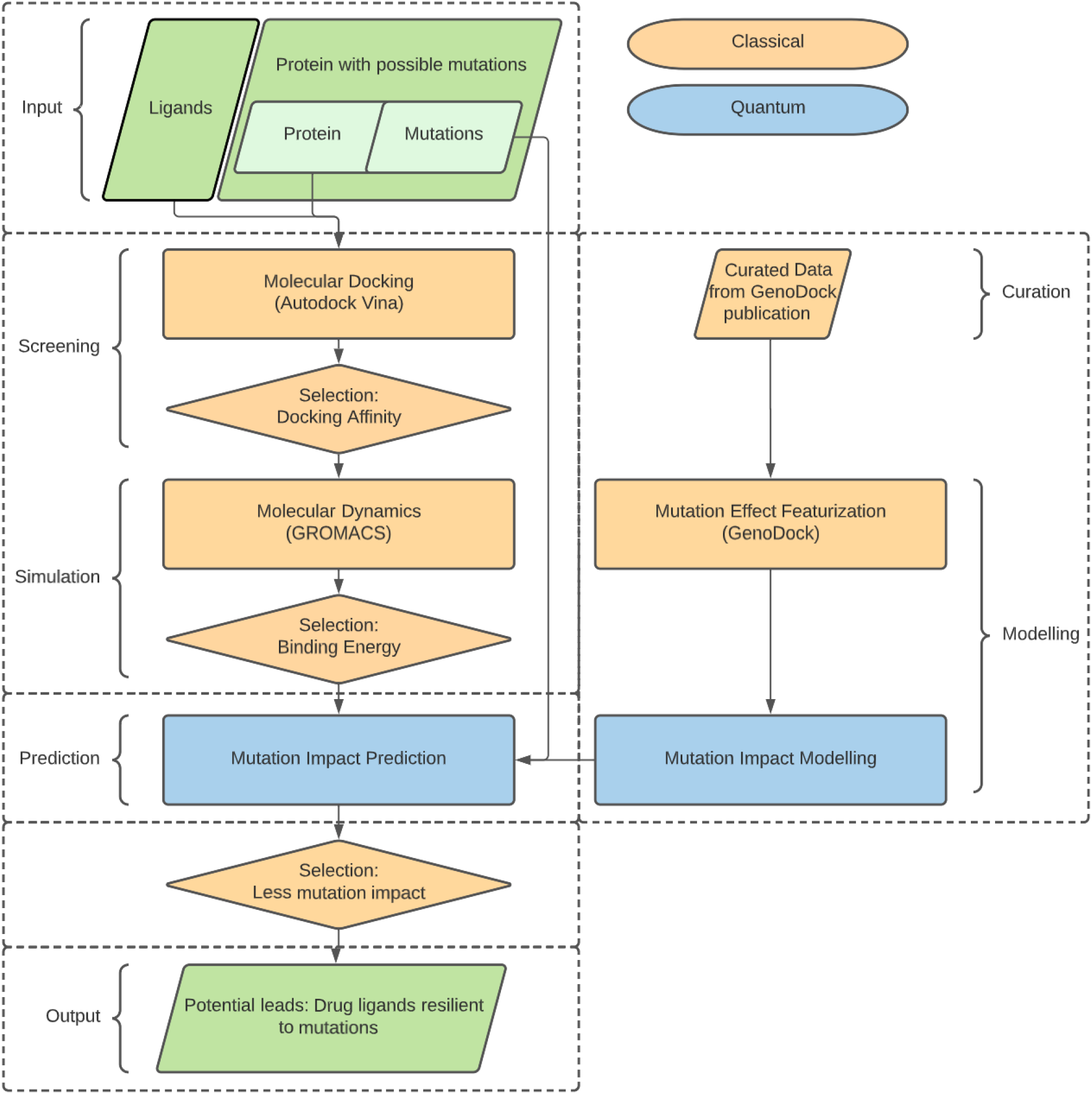
Given a set of ligands and a target protein with known or hypothetical mutations, the classical/quantum hybrid screening pipeline seeks for potential leads – ligands that would bind favorably to the protein as well as its mutants (green parallelograms, representing the input data and output results). The current implementation performs classical computation with down-selection after screening and simulation. Using a set of curated data, featurization is performed before modeling by quantum machine learning (blue rectangle). The impact of mutations on the down-selected ligands is then by the quantum model (blue rectangle). Further down-selection based on mutation impact is performed classically. We note that the selection is effectively an optimization or ranking problem, along with docking and molecular simulation, could be considered to have a quantum computational approach in the future.

This is a measured approach to the introduction of QC into the field of CADD. We prioritized the need for generating a pipeline immediately applicable to large-scale, real-world data for drug repurposing over the ultimate goal of fully exploiting putative advantages of QC. Using this work as a foundation, the QML models can be made increasingly sophisticated and eventually provide improved computational speedup and representation of structure in the data. Indeed, other modules in the docking/MD pipeline may be more inherently “quantum mechanical” in nature, as well as more computationally intensive (and therefore more in need of QC speedup). One of the main focuses of current efforts in this direction is to improve quantum chemistry simulations, which would, in turn, improve the parameterization of force-fields, and also the *ab initio* quantum mechanical calculations of binding affinity (Cao, 2018). However, some of those advances will have to wait for both hardware growth and methodological developments to allow for high-throughput screening: the number of qubits required would depend on the size of the ligand and target binding pocket, and the hybrid classical-quantum frameworks would need to be carefully designed for near-term, noisy devices. Our employment of QML has the additional advantage that ML approaches, being agnostic to the underlying causal mechanisms, are often easily generalizable to other problems (see discussion in (Cao, 2018)). For example, the numerical force-field fitting procedures in the design of programs such as AutoDock Vina (Trott et. al., 2010; Morris et. al., 2009) involve optimization of parameters to minimize the deviation of predictions from known experimental structures. This can potentially be cast as a QML problem. We hope this generality can also be exploited by the CADD community. We further emphasize, however, that our goal here is not to definitively prove the quantum advantage of these models. The exact nature and degree of putative advantages in representation and/or speedup are actively being debated by the scientific community. Our analysis is intended to lay out a clear path to QC incorporation, in the near-term and for high-throughput applications.

HypaCADD is a general-purpose workflow, applicable to any set of input drug ligands, target proteins and point mutations of the proteins. Here, we apply HypaCADD to the highly relevant problem of finding drug ligands that bind with high affinity to SARS-CoV-2 virus proteins, and whose binding is resilient against a spectrum of mutations in the target protein. Because of its importance to protein processing and viral replication, the main SARS-CoV-2 protease 3CL^pro^ has been frequently targeted (Jang et. al. 2021) for drug-ligand analyses. We use this protein and a set of associated amino acid mutations for our analyses. We initially applied a sequential virtual screening protocol, including molecular docking and molecular dynamics simulations, to ∼30,000 ligands and identified 2 potential drug candidates for the SARS-CoV-2 3CL^pro^ protein. Subsequently, we predict the effect of protein mutations that can guide lead search and optimization. Our high-throughput screening analysis will complement existing models that incorporate dynamic and structural information to assess the impact of variants on proteins and protein-ligand complexes (Torrens-Fontanals, 2021). We believe our approach increases the robustness of anti-SARS-CoV-2 drug discovery.

## Methods

### Datasets

The drug dataset (features: 3D, Ref/Mid pHs, Drug-like, In-stock) from ZINC in PDBQT format was downloaded from https://zinc.docking.org/, comprising a total of more than 10 million compounds. The X-ray crystallized 3D structure of 3CL^pro^ of HCoV-229E (PDB code: 2ZU2) (Lee et. al., 2009) and 3CL^pro^ of SARS-CoV-2 of COVID-19 (PDB code 6W63) (Mesecar, 2020) were downloaded from https://www.rcsb.org/. 2ZU2 was used as the receptor to validate our screening methodology in HCoV-229E and 6W63 was used as the receptor to identify lead compounds for SARS-CoV-2. Both receptor files were downloaded in the PDB format. Real mutations in SARS-COV-2 on nsp5 3CL^pro^ were collected from https://covidcg.org/.

### Molecular docking

AutoDock Vina (Trott et. al., 2010) was chosen as the docking tool to conduct the initial virtual screening by estimating the noncovalent binding of receptors and ligands. The receptors were processed using a custom script, removing “HETATM” components including ligands, ions and waters and protonating His41 to the neutral state at the epsilon nitrogen (Nε2). Then the preparation of a receptor for docking was finished by a script in AutoDock Tools (Morris et. al., 2009) (“prepare_receptor4.py”), where the polar hydrogens and Gasteiger charges were added and where the PDB files were converted to PDBQT format. The grid boxes for docking were centered on the active His41 residue and were set to extend 26 grid points in each direction. The lower the predicted binding affinity score is, the higher the binding affinity between the receptors and ligands.

### Molecular dynamics simulation

Molecular dynamics were performed using a combination of GROMACS (Berendsen et. al., 1995; Lindahl et. al., 2001; Van Der Spoel et. al. 2005; Hess et. al., 2008; Pronk et. al., 2013; Páll et al., 2015; Abraham et. al., 2015), AMBER03 force field (Duan et. al. 2003; Sorin et. al. 2005; Bremer et. al., 2020; Alvarado et. al., 2020), Open Babel (O’Boyle et. al., 2011), and ACPYPE (Sousa da Silva et. al., 2012; Batista et. al., 2006; Wang et. al. 2004; Wang et. al. 20006) with General Amber Force Field and TIP3P model (See Supplementary Materials). Shorter 0.4ns simulations were performed for various ligands with different binding affinities estimated by AutoDock Vina (Trott et. al., 2010). Longer simulations (10ns) were performed on selected ligands, including the ligand named X77 that was co-crystallized with SARS-CoV-2 3CL^pro^. After the simulation, GROMACS’s output is analyzed with *gmx_mmPBSA* (Valdés-Tresanco et. al. 2021; Miller et. al., 2012) which uses Ambertools 2.0’s MMPBSA to calculate binding free energy using the generalized Born surface area (GBSA) method.

### Evaluation of viral mutations’ impacts on protein-drug interactions

A classical physical-statistical classifier (Wang et. al., 2019) was developed to predict the impacts of single nucleotide variants (SNVs) on protein-drug interactions. The classifier, GenoDock, uses genomic, structural, and physicochemical features. As discussed in the Supplementary Materials, this work reformulates the framework for practical treatment of non-human proteins with less annotations. The GenoDock features per mutation-ligand configuration include (feature names are italicized and in parentheses): amino acid side-chain volume change index (*volume_change_index*), polarity change index (*polarity_change_index*), distance between the mutation and drug ligand (*distance*), molecular weight (*molecular_weight*), H-bond donor (*H_bond_donor*) and acceptor counts (*H_bond_acceptor*), rotatable bond counts (*rotatable_bond*), polar surface area (*Polar_Surface_Area*), and whether the variant occurs in the ligand binding site or not (*bind_site*). For the case of human variants, we also included features that are related to the conservation of a variant and its frequency in human populations: allele frequency (*allele_freq*); SIFT (*SIFT_consequence*) (Kumar et. al., 2009); PolyPhen-2 (*PPH_consequence*) (Adzhubei et. al., 2013); and GERP scores (*gerp_score*) (Davydov et. al., 2010). While well-studied viruses such as SARS-CoV-2 potentially allow for variant conservation properties to be assessed (see variant frequencies and SIFT scores in (Dunham et. al., 2021)), we removed such features to build non-human-genome-compatible classifiers applicable to a wider variety of viruses lacking sufficient information. Furthermore, feature space reduction also facilitates downstream calculations on the limited-qubit quantum computers considered. In general, though, the methods are designed to incorporate all features given availability of sufficiently comprehensive variant databases.

In the current study, we specifically investigate nonhuman features in the interest of applying the QNN framework to the SARS-CoV-2 3CL^pro^ protein. That is, we consider the subset of all features not specific to human genomes (namely everything except the GERP, PolyPhen2, and SIFT scores, and the allele frequency). This left us with nine features (*bind_site, distance, molecular_weight, H_bond_donor, H_bond_acceptor, rotatable_bond, Polar_Surface_Area, polarity_change_index*, and *volume_change_index*), which was a tenable number for simulated QNNs. For QNNs that run on IBM’s 5-qubit systems, we subselected again to those features which are not ligand-specific, leaving us with four features: *bind_site, distance, polarity_change_index*, and *volume_change_index*. This is a 5-qubit problem (one input qubit for each feature plus an additional readout qubit) and therefore perfectly suited to our resources. It is worth noting that, in our testing on simulated devices, we found no significant performance difference between the nine-feature group and the four-feature group. We suspect this is due to the dominance of the *bind_site* feature in the prediction process, but, regardless of the cause, it reassures us that working with the four-feature group is still a meaningful problem.

We trained various models using GenoDock’s pseudo-gold training set, which was generated by applying AutoDock Vina to a large collection of co-crystal structures from the PDB, and mapping variants from germline and somatic variant databases. We then compared all methods described below (cQNN, qisQNN, weighted mQNN, and GenoDock’s original random forest method) using the Platinum test set (Pires et. al. 2015). We applied the same models to the 6W63 mutation-ligand features. Such features were computed by GenoDock’s methodology applied to the list of SARS-CoV-2 mutations, the cleaned 6W63 crystal structure, and ligand structures. Each mutation was located within the PDB structure for the target protein, and the “wild-type” PDB structure of the protein was mutated at this site using the program Modeller (Webb and Sali 2016) (resulting in a “mutant” structure).

The convention used is to set the label as “0” for non-disruptive mutations and “1” for mutations disruptive to binding. Prior to training the cQNNs and qisQNNs, we balanced the datasets with respect to the labels, and divided the resulting dataset evenly into training/validation/testing partitions. Further information on the datasets and selection of features are provided in the Supplementary Methods.

For a more careful comparison between the classical models and quantum neural networks, we also trained classical neural networks (NNs) with approximately matching numbers of parameters. These NNs, due to parameter-number restrictions, were built using scikit-learn’s multilayer perceptrons (https://scikit-learn.org/stable/modules/generated/sklearn.neural_network.MLPClassifier.html) with a single hidden layer, fully connected with the inputs. The number of nodes in this hidden layer was adjusted to approach the QNN parameter numbers, while allowing for full connections with the input layer. For example, a 3-layer quantum neural network (architecture described below) with 4 input features would have 4 × 3 = 12 free parameters. The classical NN would have 4 weight and 1 bias parameters per hidden layer node, and so we restricted the network to have 3 nodes in the single hidden layer (= 15 free parameters).

### Quantum neural network (QNN)

Several QNN models (termed cQNN, qisQNN, and weighted mQNN, varying in architectures and training) were explored to predict mutation impact on drug binding. The training set was class-imbalanced (9,611 negatives and 670 positives), and each formulation had its own approach to treating such imbalance in the training set. We compared the performance of each model on a single, independent, experimentally measured dataset.

### Weighted mQNN

The weighted mQNN is based on the margin classifier recommended by PennyLane (Schuld et. al., 2020; Bergholm et. al., 2018) which is built on top of Python’s PennyLane and PyTorch frameworks. The original method is insufficient for an imbalanced data set, and enhancement is required. In particular, we removed the random sampling of data points, and reformulated the cost function as a weighted summation of the loss function of each training point. Details are in the Supplementary Materials.

We explored the impact of the layer and margin hyperparameters by splitting the training data in half while keeping the minority-majority ratio the same. We trained the weighted mQNN on one half of the evaluation set.

### cQNN and qisQNN

The architecture implemented herein closely follows that of Farhi and Neven (FN) (Farhi et. al. 2018) (example shown in Figure 2). The main concept is that a series of features are mapped onto an equal number of input qubits, and a training dataset is used to capture structure in the data by making sure that the measurement output of the network closely matches that provided in the training data. By iteratively adjusting the free parameters in the network, we arrive at the best performing model.

**Figure 2.**
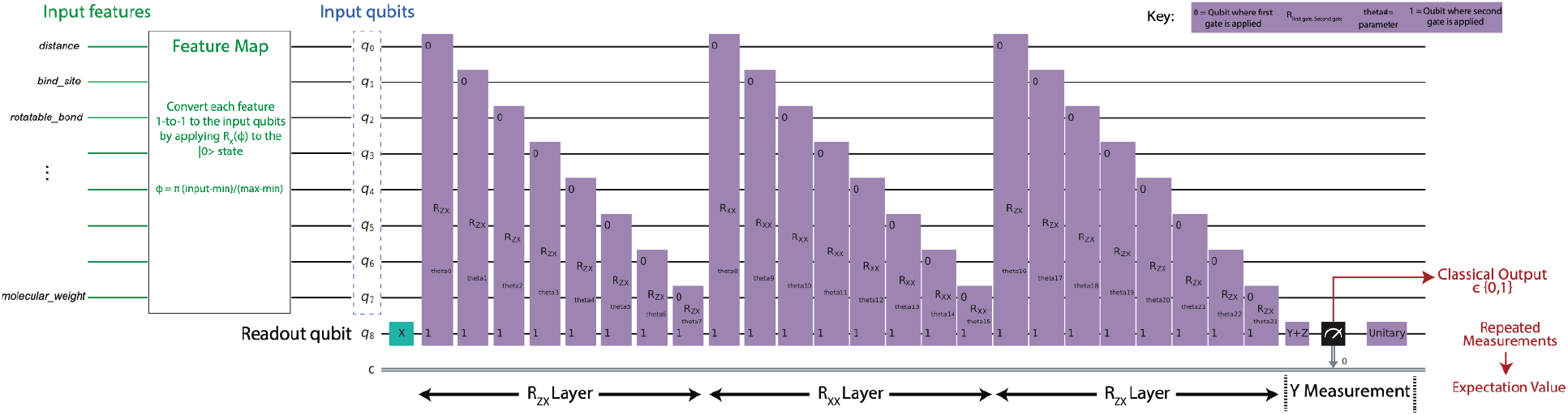
Architecture of the cQNN and qisQNN models. The number of input qubits shown here is a representative example, while in practice the number of inputs varies by the number of features considered. The initial mapping of the continuous inputs onto the interval [0, π]is not shown explicitly here.

As in FN, the QNN models here are variational quantum circuits, where the learned parameters are rotation phases applied to combinations of qubits and each circuit element takes the simple form of a unitary matrix rotation applied to a subset of the qubits: *U*(θ) = *exp*(*i*θΣ), where 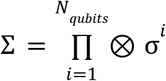 is a tensor product of operators from the set of 2 × 2 Pauli matrices {σ_*x*_, σ_*y*_, σ_*z*_} acting on a subset of the qubits. Following the simplest version of FN’s architecture, the QNN circuit consists of several *input qubits* and a single *readout qubit*. The number of input qubits equals the number of features in the dataset. The qubit z basis is assumed throughout this discussion. Each input qubit is initialized according to the data input in the following manner: (1) We directly encode binary feature values as |0 > or |1 >. (2) We apply min-max scaling, 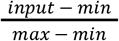, to continuous feature values for a range of [0,1]. Each input qubit is initialized according to the data via the application of an *R*_*X*_ (ϕ) operator to the |0 > state, where ϕ_*i*_ = π * *input*_*i*_ for the value of the corresponding data element. The readout qubit is always initialized to |1 >. We considered the circuit model where, across all circuit operations, input qubits only interact with the readout qubit, and not directly with each other. This choice was made purely for investigating the effect of conditioning the output directly on each input qubit, following the analysis in FN.

The interaction gates were chosen from the set of two-qubit R_ZX_(θ) and R_XX_(θ) gates, where the first subscript indicates the Pauli matrix applied to a specific input qubit and the second indicates the Pauli matrix applied to the readout qubit:

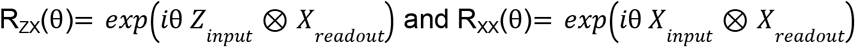

Where *Z* = σ_*z*_ and *X* = σ_*x*_, and the ⊗ symbol signifies a tensor product between the operators acting on each spin’s separate subspace (the identity operator operates on all other qubits; that is, each gate only impacts the explicitly identified qubits). A *layer* of gates is a sequence of one type of such gates applied to each of the inputs in turn.

The degree of interaction is governed by the angle θ_*n*_ of the n^th^ gate, and the θ_*n*_ are the variable parameters through which the QNN is trained. We utilized an alternating layer structure, similar to that in FN: the one-layer QNN consisted of a layer of R_ZX_ gates; the two-layer QNN of R_ZX_ followed by R_XX_ layers; the three-layer QNN of R_ZX_ - R_XX_ - R_ZX_ layers; and the six-layer QNNs involved three alternations of R_ZX_ and R_XX_ layers.

After all layers are executed, the y-component of the readout is measured, and the expectation value of that measurement is the QNN’s predicted label for the given input, 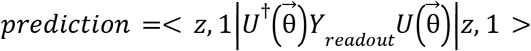, where 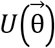 is the time-ordered product of all gates *U* (θ_*n*_) in the circuit and *Y*_*n*+1_ is the measurement of the y-component of the readout qubit. |*z*, 1 > describes input qubits initialized according to data *z* and a readout qubit initialized to |1 >. Since quantum measurement collapses the state into a single value, repeated measurements (“shots”) are required to approximate the circuit’s expectation value, given as 0 × *Fraction of zero measurements* + 1 × *Fraction of one measurements*. In our implementation, the number of circuit executions is governed by the *shots* hyperparameter. Though a higher number of shots corresponds to greater measurement accuracy, it comes at the cost of requiring more circuit executions during model training, especially during parameter update (see below). In simulation, we tested *shots* values ranging from 20 to 200 for the simulated results and found *shots*=100 to be sufficiently accurate without dramatically increasing training time. The loss function, following FN, defined as 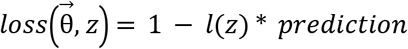, where *l*(*z*) is the true output label of the training set. This function is summed over all the data points *z* and minimized with respect to the parameters 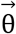. In prediction tasks, the label is assigned to 0 if *prediction* <0. 5 and 1 otherwise.

As mentioned earlier, the dataset was balanced by taking the smaller number of “disruptive” data points (i.e. mutation-ligand pairs for which AutoDock Vina indicated a positive change in the binding affinity upon mutation) and randomly selecting an equal number of non-disruptive data points. This balanced dataset was then split in a 1:1:1 ratio between training/validation/testing partitions. The training of the QNNs was carried out through batch stochastic gradient descent, where batches of 20 data points were input into the circuit, and the parameter values were updated after each such batch input. A full run-through of all data points in the training set constituted a complete training *epoch*. The number of epochs in the training depended on the performance, after each epoch, on the validation set. In the absence of improvement of validation loss after a set number of epochs (the *patience* hyperparameter), the training procedure was stopped. We found that unlike the aforementioned original mQNN, which empirically required weighted mQNN modification for good convergence (Figure 8), simple batch stochastic gradient descent was sufficient here.

We have implemented two types of variational QNNs: Models labeled “cQNN” (custom QNN) are built using custom wrapper functions that we defined using PyTorch in accordance with loss functions and a finite-difference-based parameter update defined by FN. Models labeled “qisQNN” rely on the Qiskit’s NeuralNetworkClassifier class, which uses an L1 loss function and where the parameter update is governed by a Constrained Optimization BY Linear Approximation (COBYLA) optimizer (https://qiskit.org/documentation/machine-learning/stubs/qiskit_machine_learning.algorithms.NeuralNetworkClassifier.html). The training of these models is carried out using Qiskit’s the QASM simulator (https://qiskit.org/documentation/stubs/qiskit.circuit.QuantumCircuit.qasm.html), while a limited set of test predictions based on these trained models were run on IBM Quantum’s open access 5-qubit systems (https://quantum-computing.ibm.com) (feature groups were restricted to 4 features for these test runs), specifically the *ibmq_bogota* system. For the QASM simulator, no additional noise was added, and so the only noise source was the randomness of the individual shots. We avoided adding any noise to obtain the ideal performance of the QNNs on simulators and to enable a one-to-one comparison with the noise-free classical models. On the other hand, the prediction runs on the real quantum devices were run with 1000 shots each to ensure robust statistics amid the inherent hardware noise in these devices.

To clarify the ensuing results, we emphasize that we have run many versions of these architectures with a differing number of layers, and with differing numbers of features. The variation in the layers is done to meet the restrictions on the IBM Quantum devices, and we state so in the Results section. The variation in the number of features was done to reflect different application scenarios (human vs. non-human), as described above, or to reduce the number of features down to a tractable number for the IBM Quantum devices.

Further details on the QNN training and evaluation are provided in the Supplementary Methods.

## Results

### The Workflow

We present a hybrid classical/quantum computational workflow, HypaCADD, which screens for ligands (a) that would bind favorably to a wildtype protein and (b) that would be less likely to be affected by amino acid mutations of such a wildtype protein. Because the number of potential ligands and the number of potential mutations are both large, the number of potential ligand-protein-mutation combinations quickly becomes intractable for detailed experimental follow-up. Figure 1 illustrates the multi-step screening process which uses docking and molecular dynamics to select a smaller number of candidates that would bind to a wild-type protein. Using the thus-selected candidates, we predict the impact of potential mutation with quantum machine learning after featurization. Given a combination of ligand, unmutated protein structure, and amino acid mutation, the machine learning featurization for predicting the impact of mutation is illustrated in Figure 3. The net result is a set of ligands that would bind to a target protein, and be robust to an input set of mutations.

**Figure 3.**
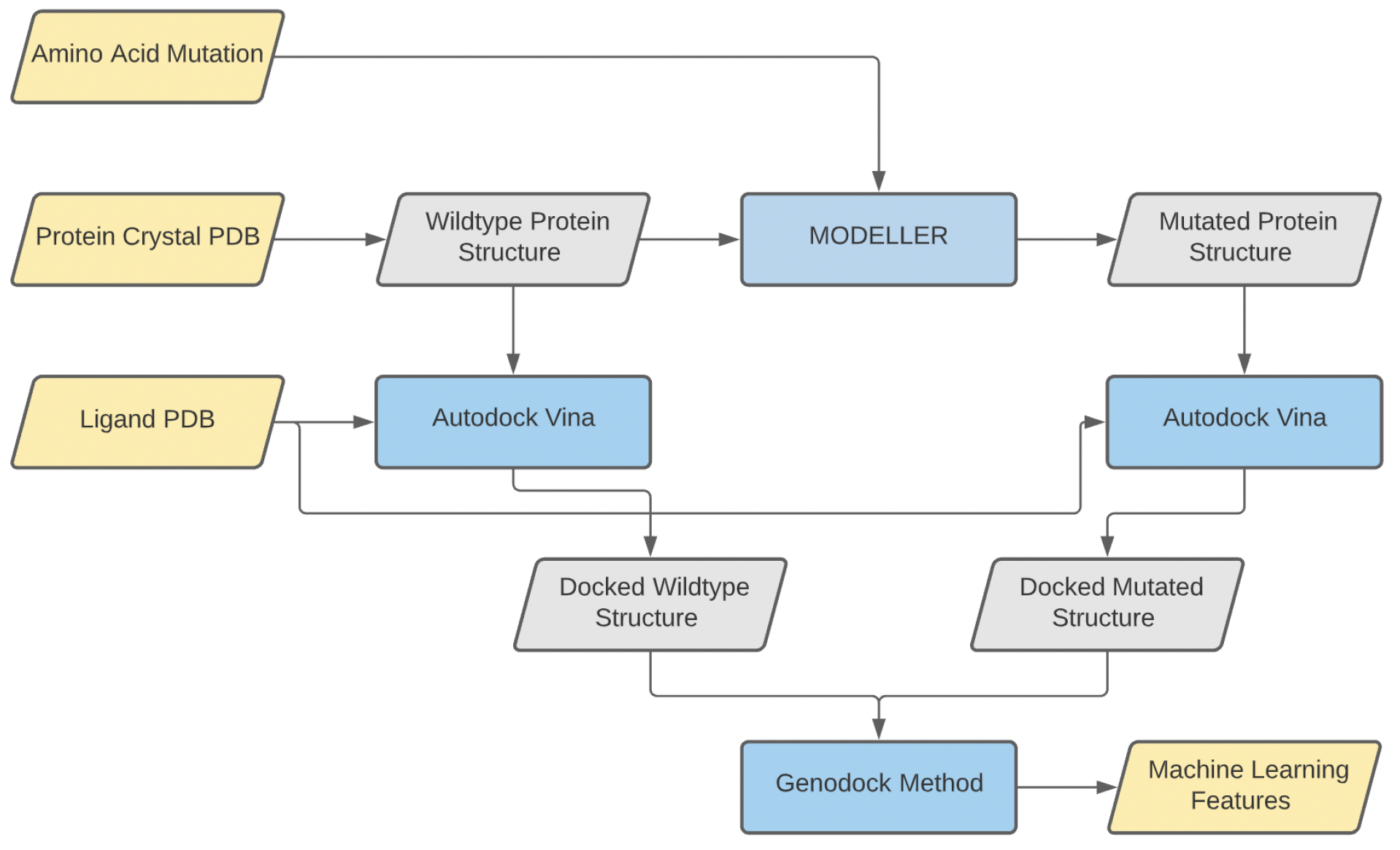
Given a combination of ligand, protein, and amino acid mutation, the features used for machine learning are computed by comparing the docked wildtype structure and docked mutated structure computed using MODELLER and AutoDock Vina. Input and output data are represented in yellow, structure data are in gray, and computational methods are in blue.

### Method validation

The paper, “Potential Broad Spectrum Inhibitors of the Coronavirus 3CL^pro^: A Virtual Screening and Structure-Based Drug Design Study,” (Berry et. al. 2015) has demonstrated an established approach to identify lead compounds, which have shown promise as inhibitors of 3CL^pro^ in coronavirus. To obtain confidence in our screening method, we validated our performance with 13 out of their 19 reported ligands which had identical ZINC IDs in the version of our drug dataset. Beside the 13 ligands, 29,981 randomly selected ligands from our drug dataset were also used as background for method validation.

The same molecular docking method, AutoDock Vina, was applied to the 13 ligands from (Berry et. al. 2015) and the randomly selected ligands, by docking against the 2ZU2 receptor, a 3D structure of 3CL^pro^ in HCoV-229E. All the 13 reported ligands had binding affinities less than -7 kcal/mol and 8 of them had binding affinities less than -9.5 kcal/mol (Table 1), which was the screening criterion used in (Berry et. al. 2015). Compared to the random ligands, the reported 13 ligands were highly enriched with ligands that can bind to the 2ZU2 receptor with high affinity (Fisher’s Exact test p-value < 2.2e-16) (Table 2), indicating we were able to identify high-quality candidates in our initial screening for potential lead compounds.

**Table 1.**
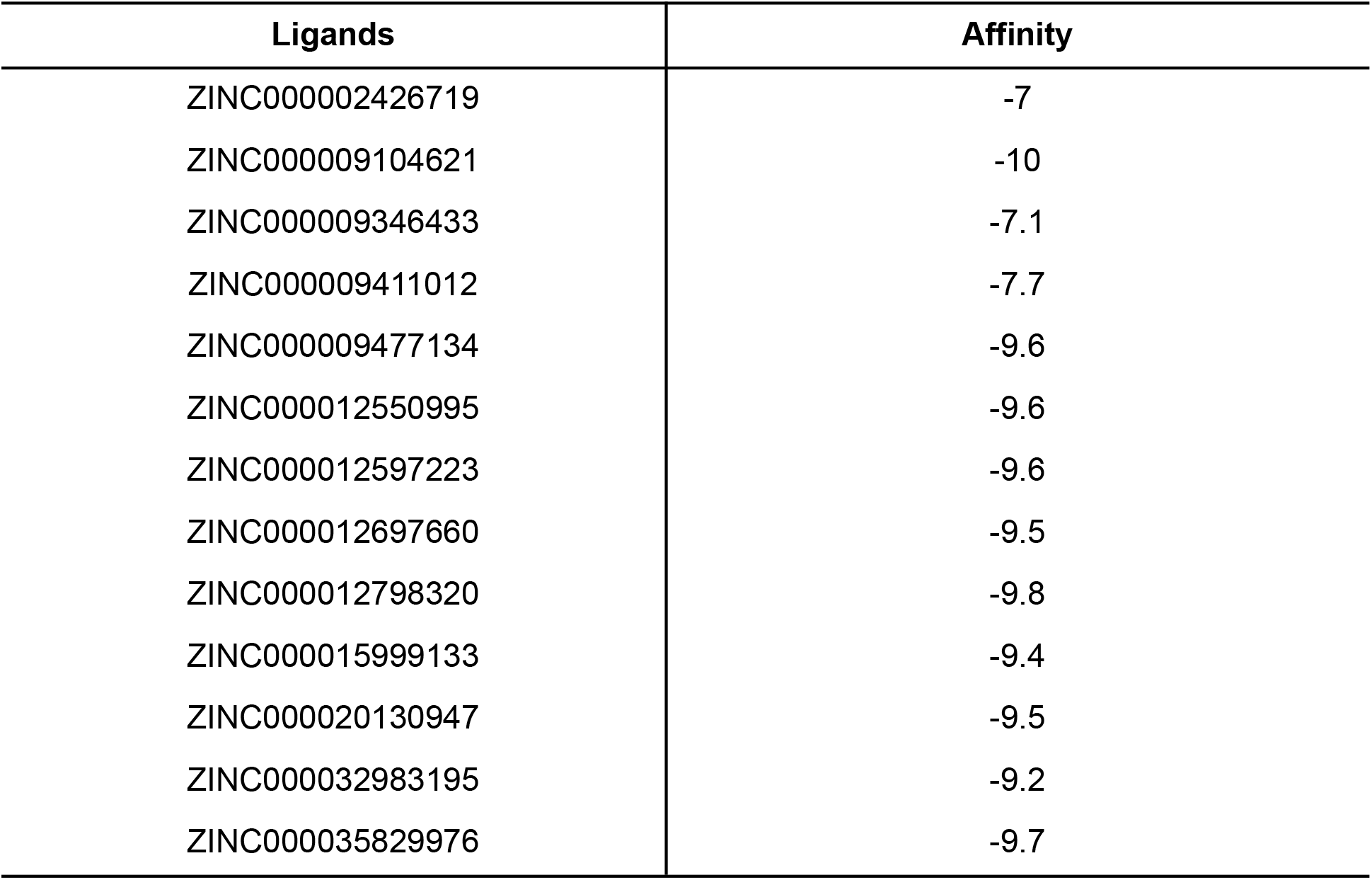
Binding affinities from molecular docking using the 2ZU2 receptor

**Table 2.** The number of ligands satisfying binding affinity cutoffs among 13 ligands from (Berry et. al. 2015) and 29,981 randomly selected ligands when docking to the 2ZU2 receptor

| Affinity criteria | (Berry et. al. 2015) | Random |
| --- | --- | --- |
| >-9.5 | 5 | 29,970 |
| ≤-9.5 | 8 | 11 |

Fisher's Exact Test: p-value < 2.2e-16

MD simulation is a more physically realistic but computationally intensive method to assess the free energy of binding between receptors and ligands. We used the MM-GBSA calculation implemented by the GROMACS software for our MD simulations. In a 0.4ns simulation, MM-GBSA was applied to a manageable subset of the aforementioned ligands with varying binding affinities predicted by AutoDock Vina. Specifically, based on Table 2, 32 ligands were selected which included all the 19 ligands having binding affinities ≤-9.5 kcal/mol and 13 ligands having binding affinities >-9.5 (with 8 selected from the random set). They were processed with MM-GBSA, as shown in Table 3. Our comparison showed that ligands with MM-GBSA scores less than -30 kcal/mol had an average of -9.5 kcal/mol AutoDock affinity score, which was significantly lower than the rest which had an average of -7.3 kcal/mol (p-value: 5.04E-04). Further, we observe that the MD GBSA binding energy of all of (Berry et. al. 2015) ligands are significantly less from the aforementioned 8 randomly drawn ligands with affinity >-9.5 (p-value: 2.7E-3, Wilcoxon rank-sum). While the Pearson correlation of AutoDock affinity and MM-GBSA was 0.668, it demonstrated that predictions from AutoDock were indicative but the accuracy can largely increase with more sophisticated MD simulations.

**Table 3.** Free energy of binding to the 2ZU2 receptor predicted by MM-GBSA for a collection of 32 ligands, with 19 of them having an affinity ≤-9.5 kcal/mol and 13 of them >-9.5 kcal/mol. Note: Ligands ZINC000020130947 and ZINC000002467880 failed in MD simulation.

| Ligand | Affinity | GBSA |
| --- | --- | --- |
| ZINC000002426719 | -7 | -30.7826 |
| ZINC000002473650 | -5.6 | -24.1817 |
| ZINC000002484659 | -4.8 | -12.0924 |
| ZINC000004533051 | -5.2 | -25.4426 |
| ZINC000005121078 | -5.3 | -24.2796 |
| ZINC000008386858 | -5.7 | -23.3873 |
| ZINC000009104621 | -10 | -36.9424 |
| ZINC000009346433 | -7.1 | -27.0533 |
| ZINC000009411012 | -7.8 | -22.1088 |
| ZINC000009477134 | -9.6 | -33.2358 |
| ZINC000009518465 | -10.1 | -37.5634 |
| ZINC000009731745 | -9.7 | -29.1525 |
| ZINC000012550995 | -9.6 | -30.9933 |
| ZINC000012597223 | -9.6 | -43.1545 |
| ZINC000012697660 | -8.6 | -21.0867 |
| ZINC000012798320 | -9.8 | -34.5397 |
| ZINC000015953686 | -9.4 | -28.8171 |
| ZINC000015959761 | -9.7 | -35.7219 |
| ZINC000015999133 | -9.4 | -47.1014 |
| ZINC000016001299 | -9.6 | -26.5863 |
| ZINC000016020583 | -10.2 | -22.4507 |
| ZINC000020533854 | -9.9 | -40.7371 |
| ZINC000032983195 | -9.3 | -39.0102 |
| ZINC000035829976 | -9.7 | -27.0573 |
| ZINC000057129043 | -9.6 | -33.8203 |
| ZINC000088375393 | -5.5 | -9.9067 |
| ZINC000101766867 | -9.4 | -46.0811 |
| ZINC000225765471 | -5.9 | -24.3948 |
| ZINC000247300899 | -9.5 | -45.4873 |
| ZINC000386725971 | -6.1 | -24.3231 |

Overall, our results demonstrated that virtual screening and simulation combined can reliably serve as the foundation for predicting lead compounds for further optimization and selection.

### Defining benchmarks and accuracy of virtual screening for SARS-CoV-2 3CL^pro^

Recently, there have been several resolved X-ray crystallographic structures of SARS-CoV-2 3CL^pro^; however, most of those structures are complexed with an irreversible substrate-like inhibitor. At the time of this study, only one structure (PDB code: 6W63) represented SARS-CoV-2 3CL^pro^ with a reversible dipeptide inhibitor (X77). This structure and inhibitor could serve as a benchmark for virtual drug screening for SARS-CoV-2 3CL^pro^. To validate the docking method for SARS-CoV-2 3CL^pro^, AutoDock Vina and GROMACS MD were applied to the receptor and inhibitor X77 in the 6W63 structure. The root-mean square deviation (RMSD) between the experimental structure of X77 and best predicted pose of docking by AutoDock Vina was 0.814 Å, which was below the well-defined 2 Å benchmark to assess the accuracy of docking methodology (Wagner et. al. 2019). The overlay of the crystal structure of X77 and docked pose is shown in Figure 4. The binding affinity and MM-GBSA between the receptor and inhibitor of 6W63 were -8.3 kcal/mol and -37.69 kcal/mol, respectively.

**Figure 4.**
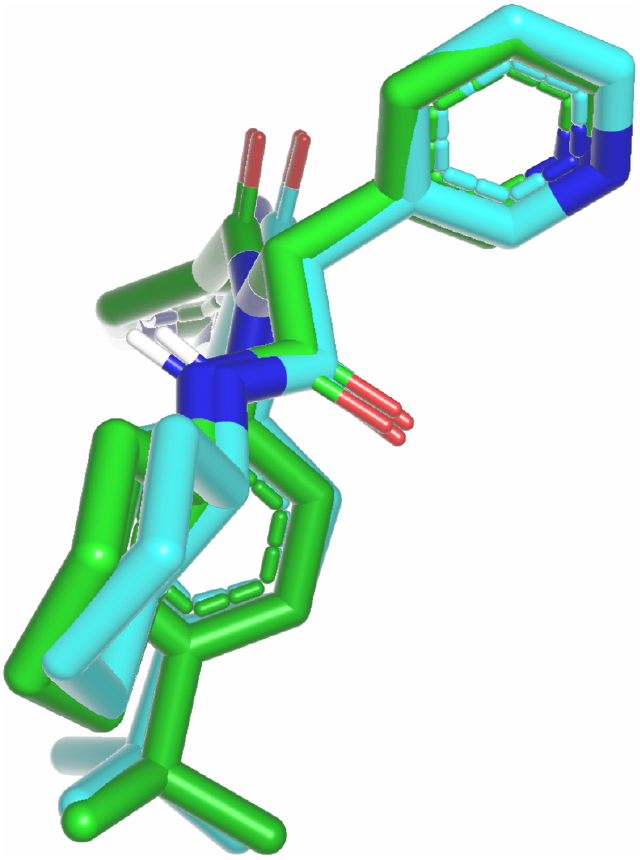
The overlap of the crystal structure of X77 and docked pose.

### Molecular docking and molecular dynamics for SARS-CoV-2 3CL^pro^

Confirming results of (Berry et. al. 2015) and benchmarking on the 6W63 structure provided us confidence to use AutoDock Vina and GROMACS MD as high throughput tools for virtual screening. We further applied our protocol to SARS-CoV-2 3CL^pro^ against the 29,981 randomly selected ligands and 13 previously reported ligands, as aforementioned. Comparison of AutoDock results of the two receptors showed the ligands’ binding affinities to 2ZU2 and 6W63 were highly correlated but some differences existed as shown in Figure 5 and S4, which might be due to the highly similar but not identical amino acid sequences between HCoV-229E 3CL^pro^ and SARS-CoV-2 3CL^pro^. For example, the 13 ligands highlighted in Figure 5 shows that ligands with a high affinity (<-9.0) for 2ZU2 did not have affinities as high for 6W63. There were 40 ligands (Table 4) with binding affinity ≤ -9.5 kcal/mol for SARS-CoV-2 3CL^pro^, of which only 4 overlapped with those for HCoV-229E (Supplementary Figure S4).

**Figure 5.**
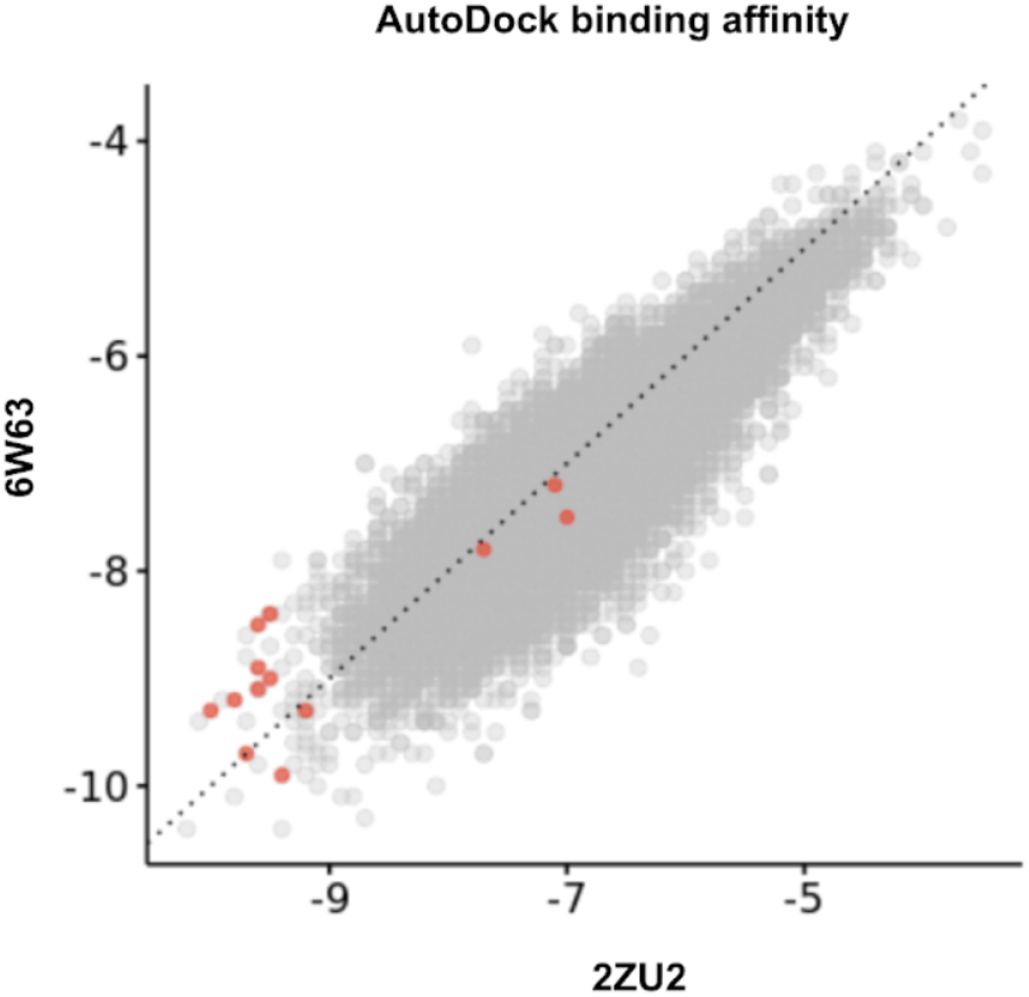
Comparison of AutoDock binding affinity. The scatter plot of AutoDock binding affinity between the 2ZU2 and 6W63 receptors. The highlighted dots represent the 13 reported ligands.

**Table 4.**
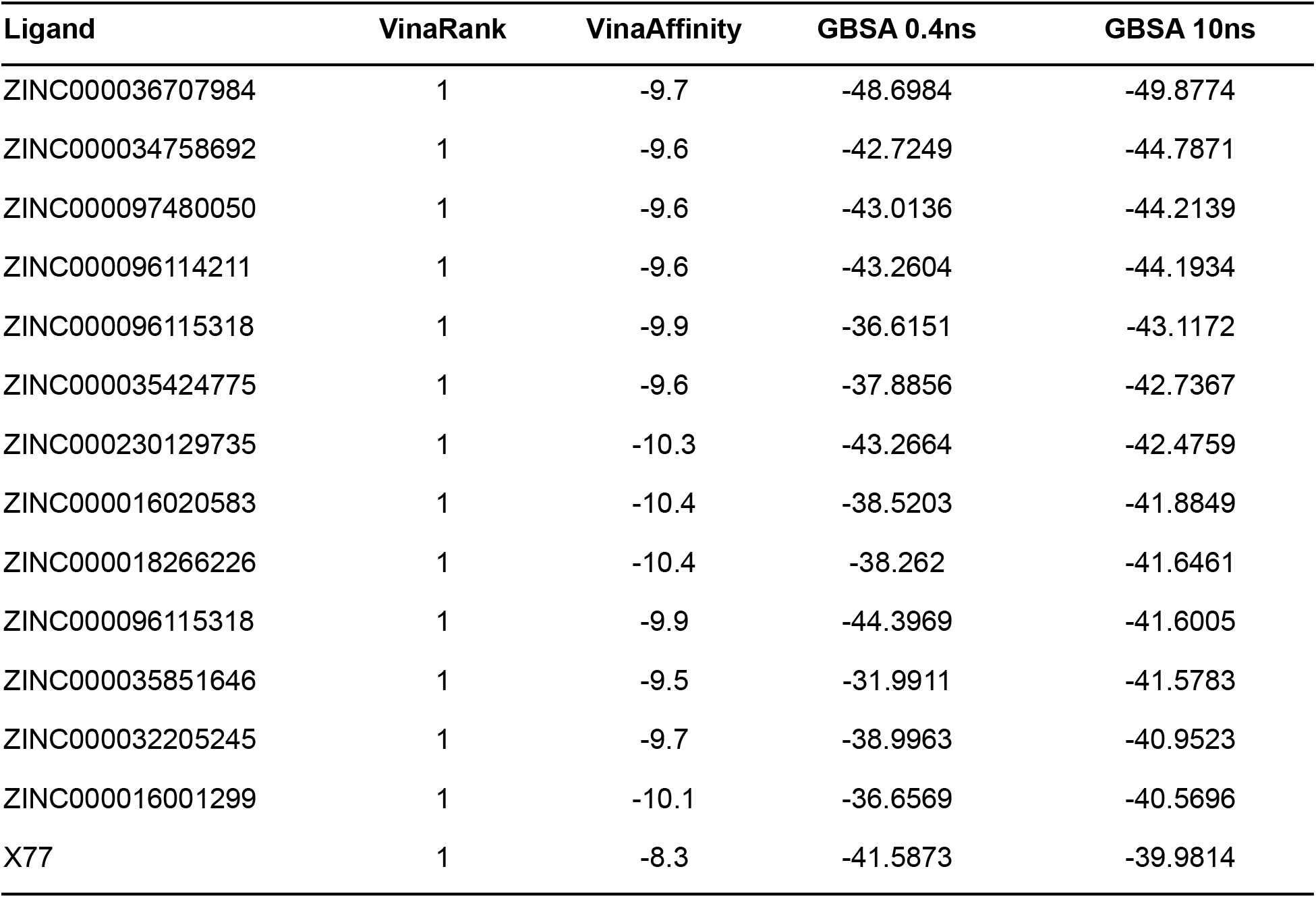
Binding affinities (≤-9.5 kcal/mol) from molecular docking using the 6W63 receptor and free energies of binding predicted by MM-GBSA that were lower than that of X77 in a 10ns MD simulation. See Supplementary Materials for the full table of the 40 ligands.

| Ligand | VinaRank | VinaAffinity | GBSA 0.4ns | GBSA 10ns |
| --- | --- | --- | --- | --- |
| ZINC000036707984 | 1 | -9.7 | -48.6984 | -49.8774 |
| ZINC000034758692 | 1 | -9.6 | -42.7249 | -44.7871 |
| ZINC000097480050 | 1 | -9.6 | -43.0136 | -44.2139 |
| ZINC000096114211 | 1 | -9.6 | -43.2604 | -44.1934 |
| ZINC000096115318 | 1 | -9.9 | -36.6151 | -43.1172 |
| ZINC000035424775 | 1 | -9.6 | -37.8856 | -42.7367 |
| ZINC000230129735 | 1 | -10.3 | -43.2664 | -42.4759 |
| ZINC000016020583 | 1 | -10.4 | -38.5203 | -41.8849 |
| ZINC000018266226 | 1 | -10.4 | -38.262 | -41.6461 |
| ZINC000096115318 | 1 | -9.9 | -44.3969 | -41.6005 |
| ZINC000035851646 | 1 | -9.5 | -31.9911 | -41.5783 |
| ZINC000032205245 | 1 | -9.7 | -38.9963 | -40.9523 |
| ZINC000016001299 | 1 | -10.1 | -36.6569 | -40.5696 |
| X77 | 1 | -8.3 | -41.5873 | -39.9814 |

Furthermore, our MD simulations ran successfully on all the 40 ligands having binding affinity ≤ -9.5 kcal/mol and on X77 (Table 4). We surmised that a longer simulation time, such as 10ns, would yield more stable and accurate results (though it takes a much longer runtime) and so we performed a 10ns simulation in addition to the 0.4ns to achieve the best results for 6W63. We observed a Spearman correlation of 0.696 (p-value=2.16E-7) between the 0.4ns and 10ns simulations. Results in Table 4 show that there were 13 ligands in the 10ns simulation with a GBSA score lower than that of X77. Of these ligands, ZINC000036707984 had the lowest GBSA and ZINC000016020583 had the lowest Vina Affinity, indicating these two ligands could have a higher or comparable binding affinity to the 6W63 receptor when compared to X77. Therefore, they were selected as a manageable set of potential lead compounds for our further investigation. Along with the reference inhibitor X77, they were evaluated for the potential impacts resulting from possible variations in the 6W63 receptor.

### Evaluation of virus mutations’ impact on the interactions of the lead compounds with SARS-CoV-2 3CL^pro^

GenoDock (Wang et. al. 2019) is a hybrid physical-statistical classifier to predict the impacts of variants on protein-drug interactions using genomic, structural, and chemical features; however, due to the focus on human proteins, some of the features used in the published GenoDock classifier were human-specific and not applicable to viral genomes. In order to make the GenoDock framework applicable to non-human genomes such as viruses and to further improve the predictive performance, we made several modifications and improvements including removing human-specific features (all the conservation scores and allele frequencies), feature normalization, and making use of an ensemble method. The final set of features employed are: volume change, polarity change, distance, molecular weight, H-bond donor, H-bond acceptor, rotatable bond, polar surface area, bind site True/False. Table 5 demonstrates that the new virus-genome compatible classifier with Random Forest implementation had comparable performance compared to the original GenoDock classifier with Random Forest implementation based on a 10-fold cross-validation.

**Table 5.** Comparison of the original GenoDock classifier and new classifier, which is virus-genome compatible after removing human-specific features. Both classifiers were implemented using the Random Forest machine learning model. The numbers in parenthesis are standard deviations of the performance scores.

|  | Original | Human-specific features removed |
| --- | --- | --- |
| **Accuracy** | 0.965 (0.005) | 0.964 (0.006) |
| **F1** | 0.701 (0.040) | 0.705 (0.047) |
| **recall** | 0.640 (0.050) | 0.663 (0.061) |
| **precision** | 0.779 (0.054) | 0.756 (0.056) |

### Classical Computing model performance on external validation set

We utilized Genodock’s 10,281 “pseudo-gold-standard” training points based on the use of AutoDock Vina to quantify the impacts of amino acid variations on ligand binding, as well as 86 validation data points from the independent Platinum database (Pires et. al. 2015) for our non-human-feature classifiers. The Platinum database results originate from experimental measurements of mutation effects. We retrained a model using GenoDock’s class-balanced random forest method with the pseudo-gold-standard set, and evaluated against the Platinum validation set. The classification method yielded an AUC of 0.628, which was on par with the original publication. Given the relative scarcity of experimental data measuring the impacts of amino acid mutation on ligand binding, and our consequent training on a computational binding affinity dataset, we see this result as providing reasonable validation for our method. We, however, hope that the future use of experimental data to train models would further improve the agreement.

### Quantum Computing model performance on external validation set

Multiple formulations of quantum neural networks have been explored to compare the performance and practicality of customized versus published formations. While the primary focus of this paper is on demonstrating the application of our computational workflow on SARS-CoV-2 proteins, we also trained our cQNN and qisQNN models on all the feature groups of the original GenoDock publication, including human-specific features such as conservation scores. Our results on the GenoDock and Platinum datasets (Supplementary Figure S1 and S2, respectively) indicate very similar performance of three-layer QNNs with the random forest model (in terms of AUC values). In all cases, the performance of the (approximately parameter-number-matched) classical NNs was on par with the best QNN model. With this confidence in the performance of the QNNs on a more general set of feature groups, we proceeded to apply the cQNN and qisQNN models to the nonhuman feature group relevant to the SARS-CoV-2 protein. These models performed very well on a GenoDock leave-out test set (Supplementary Figure S3). The results for the Platinum dataset below are from three-layer cQNNs and qisQNNs.

The weighted margin QNN (weighted mQNN, see Methods) is a reformulation of PennyLane’s margin classifier (unweighted and trained on 500 random samples from training set per iteration). Nonhuman features from Genodock are fit into a 4-qubit formulation. Due to the high computational cost, we trained the model with one half of the pseudo-gold-standard training set, then validated on the hold-out part of the training set. Figure 6 shows the convergence of the training process. It reveals that the weighted cost function stabilized the convergence when compared to the unweighted. The classification performance is demonstrated in Figure 7. It shows that the mQNN could achieve an AUC as high as 0.937 on the validation set.

**Figure 6.**
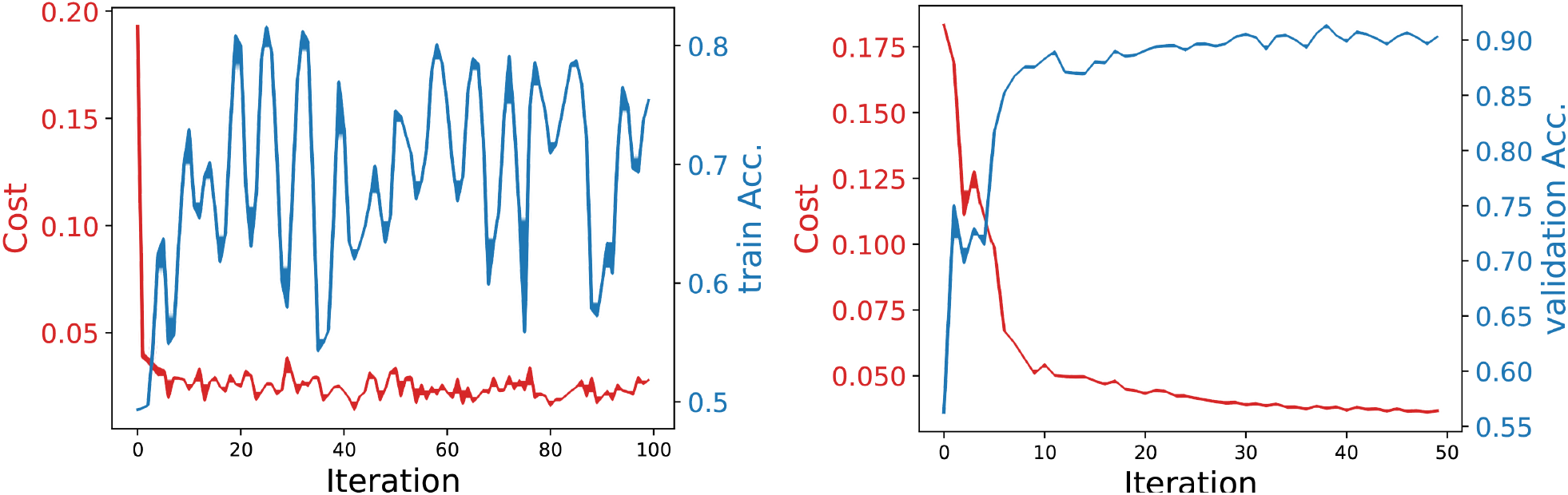
Convergence behavior (using 6 layers and a 0.15 margin) between PennyLane’s margin classifier (left) and the weighted mQNN (right). Note that the “train Acc.” is estimated by scaling up the contribution of the minority class.

**Figure 7.**
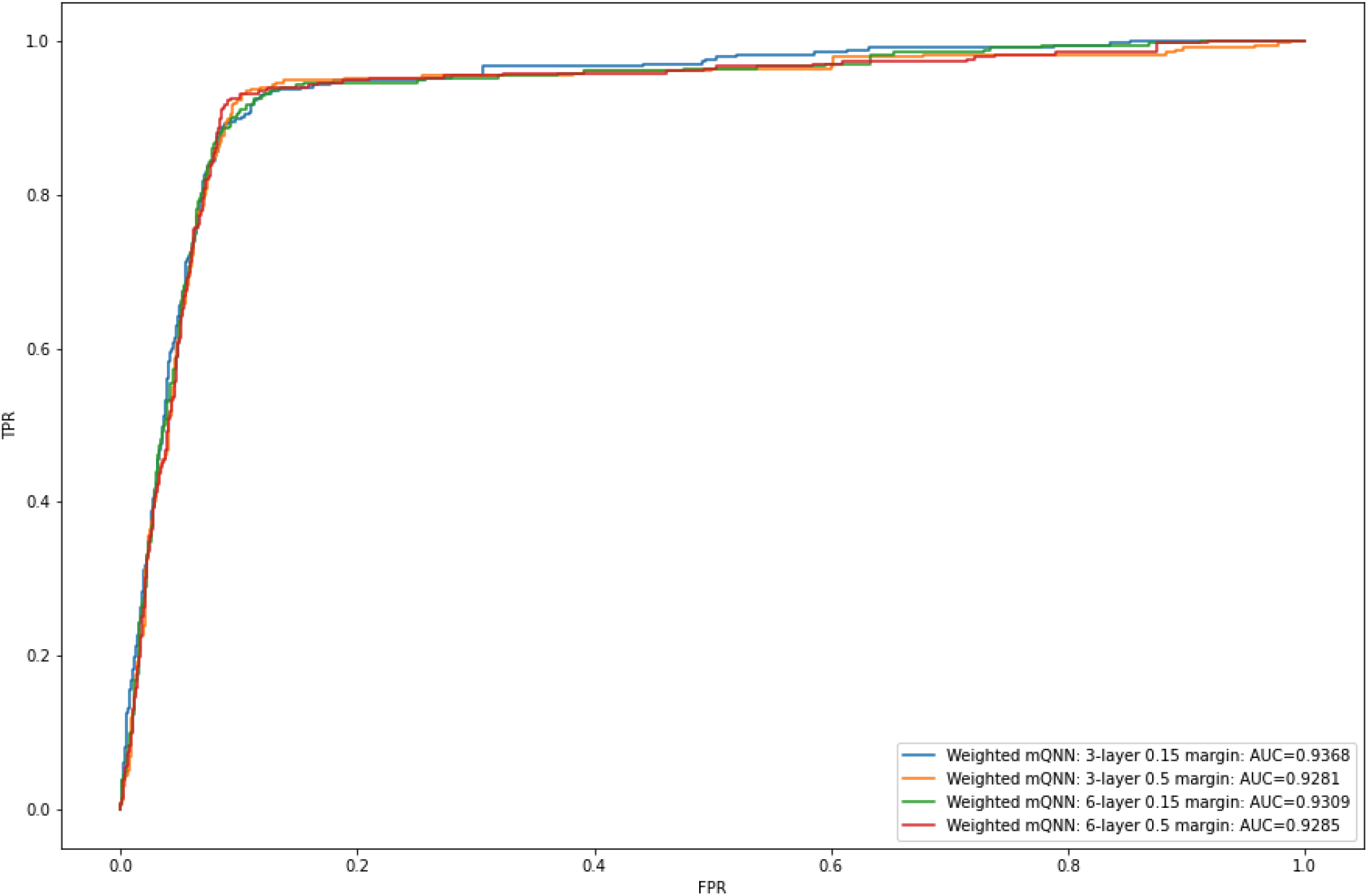
TPR vs FPR for the weighted mQNN formulation.

**Figure 8:**
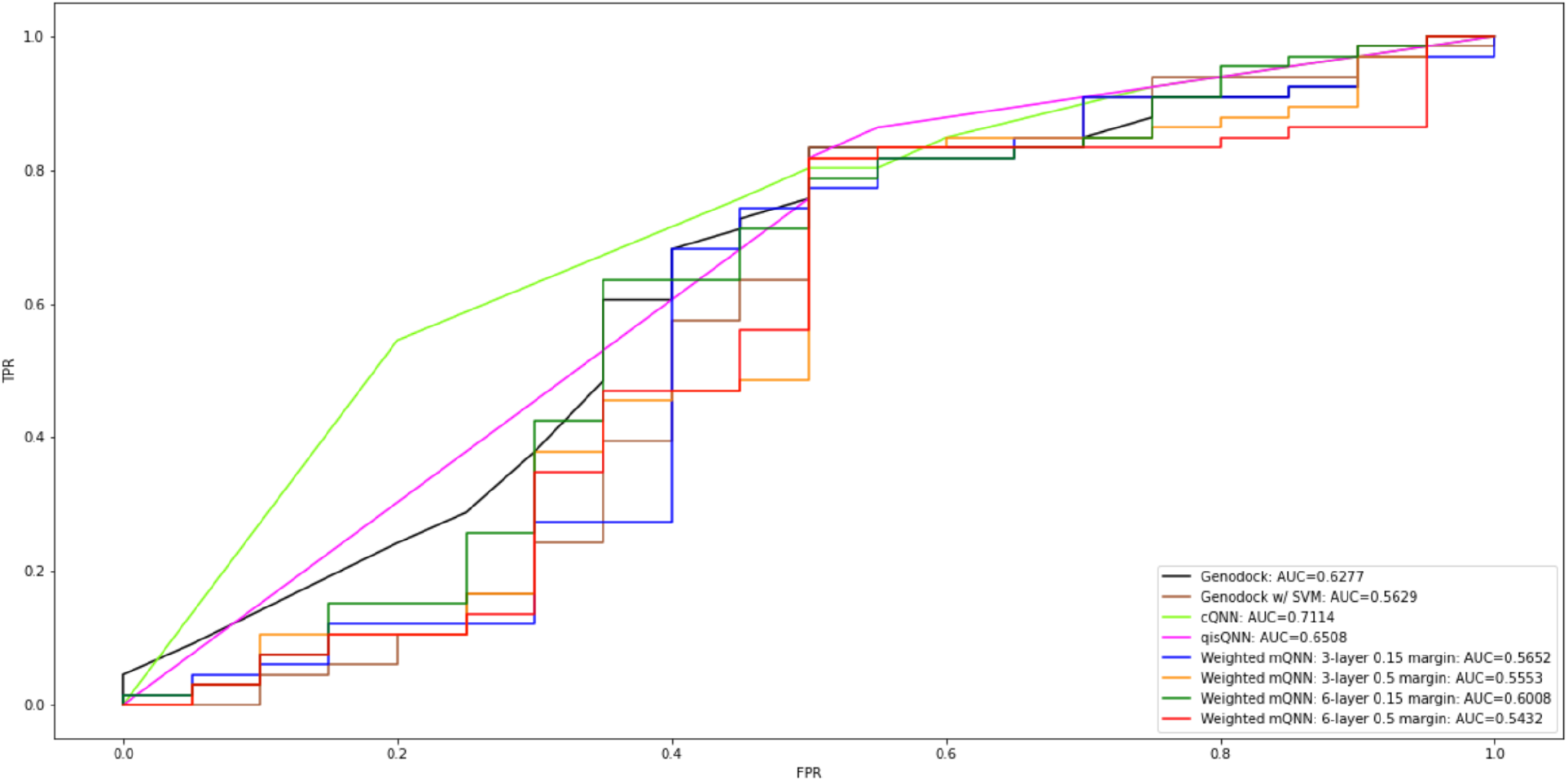
Receiver operating characteristic evaluated on the 86 validation data points from the Platinum database.

We applied our quantum modeling methods with quantum simulators to the Platinum dataset for the comparisons in Figure 8 and Table 6. Figure 8 shows the true positive rate versus false positive rate, as positive calls were made by applying different thresholds to measure, for example, the z-projection of output qubit. We note that the density of points for each curve can vary depending on the number of tie scores (Table S3). Table 6 shows the F1, sensitivity, and precision of binary prediction by the model, as well as the AUC (Supplementary Methods 3a), which is a binary-threshold-invariant measurement of model performance. The AUC indicates the sensitivity-specificity trade-offs if one wants to, for example, increase specificity at the cost of sensitivity. We also note that the precision was calculated from binary calls under heavy class imbalance in favor of positive labels, as discussed in the Supplementary Materials.

**Table 6.** Sensitivity, precision, specificity, and F1 score of the models with default classification threshold.

| Method | AUC | F1 | Sensitivity | Precision |
| --- | --- | --- | --- | --- |
| GenoDock | 0.63 | 0.83 | 0.88 | 0.79 |
| GenoDock w/ SVM | 0.56 | 0.83 | 0.82 | 0.84 |
| cQNN | 0.71 | 0.82 | 0.80 | 0.84 |
| qisQNN | 0.65 | 0.82 | 0.80 | 0.84 |
| Weighted mQNN: 3-layer, 0.15 margin | 0.57 | 0.80 | 0.77 | 0.84 |
| Weighted mQNN: 3-layer, 0.5 margin | 0.56 | 0.83 | 0.82 | 0.84 |
| Weighted mQNN: 6-layer, 0.15 margin | 0.60 | 0.82 | 0.80 | 0.83 |
| Weighted mQNN: 6-layer, 0.5 margin | 0.54 | 0.83 | 0.82 | 0.84 |

The cQNN method outperformed all other methods, classical or quantum, in terms of AUC and sensitivity in the high specificity regime. This demonstrates the value of specialized quantum neural networks. The corresponding classical NN (not included in Figure 8 for clarity) also performed well, showing an AUC of 0.68. However, we did notice that the performance of the QNN models on the external validation set was heavily correlated with a single variable: *bind_site*, a binary indicator variable labeling whether a mutation occurs within the binding site of a ligand or not. For example, if the mutation was experimentally observed to be disruptive but did not occur in the binding site, the cQNN and qisQNN models would incorrectly label the data point as non-disruptive. This was also borne out in the cross-feature group comparisons we conducted using the original GenoDock and Platinum datasets (Figures S1 and S2), where feature groups that included the *bind_site* and *distance* parameters (both of which are related) performed better. The outsize influence of these variables on the performance of the models was also found in the GenoDock analysis (Wang et. al. 2019), but does not imply the absence of influence by other features as seen in the original publication. There are some subtle differences in performance between the different feature groups that all include *bind_site* as a feature. It is likely, however, that fully capturing more complex cross-feature interactions would need more qubits and more complex gate architectures. For the mQNN, we see definite performance improvement when class imbalance is addressed by the weighted cost function. We see increased performance when the network is increased from 3 layers to 6 layers and margin is decreased from 0.5 to 0.15, and when class imbalances are compensated by weighting of the loss function.

### SARS-CoV-2 variant impact predictions in simulation and on real QC devices

With the quantum formulation established to be at least on par or better than non-quantum methods, we applied the models to predict the impact of amino acid substitution in the SARS-CoV-2 virus 3CL^pro^ on protein-ligand binding. As of Jan 2021, 98 amino acid substitution variations had been detected in SARS-CoV-2 3CL^pro^ (Figure 9). The ligand candidates were chosen to be the X77 ligand for its known crystalized structure with 6W63, ZINC000016020583 for its lowest docking affinity, and ZINC000036707984 for its lowest GBSA value (Table 4).

**Figure 9:**
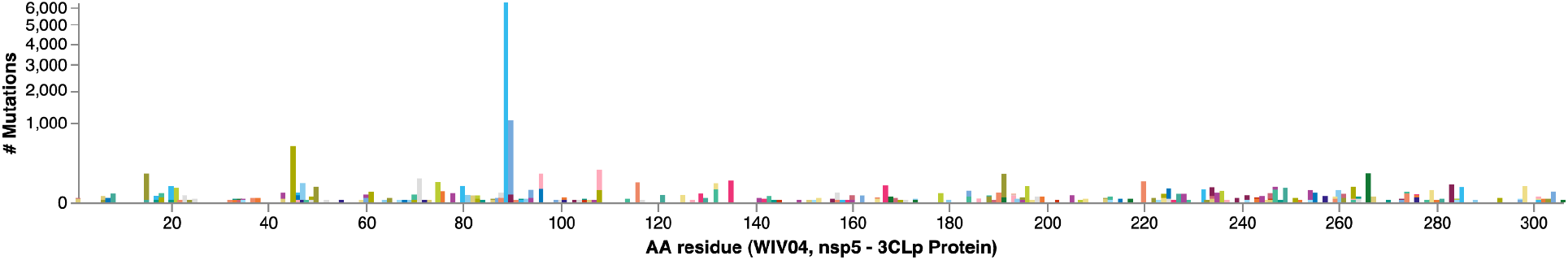
Amino acid variations distribution on SARS-CoV-2 3CL^pro^. The number of a particular variant observed in all sequenced SARS-CoV-2 samples are shown on the y-axis.

Because of the scarcity of quantum computer time, we trained all models using quantum simulators. We subsequently established consistency between test set prediction results when executed on quantum simulators and on real quantum computers. Again, addressing resource limitations, we separately trained one- and two-layer cQNNs on simulators and used those for prediction on the real devices. We selected, by Vina docking affinity change estimation, 3 mutations that would likely cover some positive and negative calls for the ligands, and compared the prediction from simulator and real quantum computers. Table 7 shows that cQNN, the best performing model, was consistent between simulation and 1000-shot measurement on a IBM 5-qubit device (the *ibmq_bogota* system) with a quantum volume of 32. Interestingly, we had to increase the number of shots relative to the simulation results in order to get robust statistics. Additionally, we found that the noise in the output expectation values increases with an increase in the number of layers, implying that each set of new gates added to a circuit brings further noise. This can be seen in Supplementary Table S1, where in addition to the same predicted output values as in Table 7, we included the expectation values output by one- and two-layer circuits (and thresholded at 0.5 to determine the final output label of 0 or 1). With 2 layers, the expectation values appear to cluster closer to the threshold of 0.5, relative to the one-layer case. In summary, while the expectation is that simulated values will deviate from real device-generated results due to decoherence and noise in the real devices, for the small number of data points considered we find no difference in predictions. However, we see the effects of device noise in the expectation values observed and in the number of shots required for robust statistics.

**Table 7.** Comparison of outputs between simulation and real quantum computer on the 3CL^pro^ mutations. Predictions are indicated as 0 or 1 depending on whether a mutation is predicted to be non-disruptive or disruptive to binding of the ligand, respectively. The results shown include: a 3-layer (Farhi and Neven architecture) cQNN circuit where both training and predictions were run using the QASM simulator; and 1- and 2-layer cQNN circuits trained on the QASM simulator, but with prediction runs on an IBM Quantum 5-qubit system (marked by the term “real”), with 1000 shots in each case; a weighted mQNN circuit with predictions conducted through simulation (default Pennylane simulator with 6-layer and 0.15 margin); a 6-layer, 0.15 margin, weighted mQNN circuit with predictions conducted through runs on an AWS’s Braket DM1 system, 1000 shots; a 6-layer, 0.15 margin, weighted mQNN circuit with predictions conducted through runs on a Rigetti system, 1000 shots.

| Position | Ref | Alt | Ligand | cQNN<br>simulation | cQNN 1-layer<br>5-qubit IBM,<br>1000 shots | cQNN 2-layer<br>5-qubit IBM,<br>1000 shots | weighted mQNN<br>6-layer, 0.15<br>margin,<br>PennyLane<br>simulation | weighted mQNN<br>6-layer, 0.15<br>margin, DM1,<br>1000 shots | weighted mQNN<br>6-layer, 0.15 margin,<br>Rigetti, 1000 shots |
| --- | --- | --- | --- | --- | --- | --- | --- | --- | --- |
| 160 | CYS | PHE | X77 | 0 | 0 | 0 | 1 | 1 | 0 |
| 168 | PRO | SER | X77 | 1 | 1 | 1 | 1 | 1 | 0 |
| 188 | ARG | SER | X77 | 1 | 1 | 1 | 1 | 1 | 0 |
| 160 | CYS | PHE | ZINC000016020583 | 0 | 0 | 0 | 1 | 1 | 1 |
| 168 | PRO | SER | ZINC000016020583 | 1 | 1 | 1 | 1 | 1 | 0 |
| 188 | ARG | SER | ZINC000016020583 | 1 | 1 | 1 | 1 | 1 | 0 |
| 160 | CYS | PHE | ZINC000036707984 | 0 | 0 | 0 | 0 | 0 | 0 |
| 168 | PRO | SER | ZINC000036707984 | 1 | 1 | 1 | 1 | 1 | 0 |
| 188 | ARG | SER | ZINC000036707984 | 1 | 1 | 1 | 1 | 1 | 0 |

We also observed that, unlike the IBM platform, the simulated values produced by PennyLane’s analytical simulator agreed poorly with those produced on the Rigetti Aspen-10 hardware. To understand this, we used the AWS’s SV1 and DM1 simulators to perform 1000-shot simulation with the PennyLane framework, and the SV1 and DM1 results agreed with those of PennyLane’s simulator. We also used the AWS’s Braket API to perform simple individual qubit RX rotations followed by z-direction measurements as well as to perform pairwise CNOT operations followed by z-direction measurements. Using the *disable_qubit_rewiring* option in the API, we disabled automatic reassignment of the API’s logical qubits onto physical qubits, and we observed that the z-direction measurements deviated from simulation/analytical values on a physical qubit-by-qubit basis, and the amount of deviation could be as much as one order of magnitude from one physical qubit to another. This qubit-dependent, heterogeneous noise could in principle be mitigated by a weighted mQNN with 4 qubits and limited nearest-neighbor CNOT operations, which maps well onto the nearest neighbor physical connectivity of the Rigetti system (e.g. qubit 1/2/15/16 in Supplementary Figure S5). However, the encoding of 10 features into 4 qubits is performed by an extra sequence of gate operations (Mottonen et. al., 2005) including additional pairwise connectivity that is not present in the Rigetti’s topology if the operations were to be restricted to involve only 4 qubits. From AWS’s reported metadata, we observed that the PennyLane 4-qubit circuit was transpiled into operations with a higher number of gates and operations before execution on the Rigetti machine. We further confirmed this by modifying the aws-pennylane plugin to disable qubit reassignment and transpilation, which induced an infrastructure error about missing 2-qubit coupling for the CNOT operations (confirmed to be the ones missing from the hardware) that were required to encode 10 features into 4 qubits. While the PennyLane framework can indeed produce consistent results among all three simulators, actual execution of non-trivial circuits on real physical hardware such as Rigetti might be hampered by qubit-dependent, heterogeneous noise. We observed that cQNN, developed within the IBM ecosystem, has the best reproducibility between simulated and real quantum simulation.

Having studied the correlations between real and simulated quantum computation, we make more predictions with the quantum simulator. The models’ predictions are listed in Figure S2. For binary prediction over all mutations and ligands, the cQNN and qisQNN had 100% concordance and the cQNN and Genodock had 99.7% concordance, whereas cQNN and weighted mQNN had 75.2% concordance. To note, X77 was predicted to be susceptible to the mutations VAL20ILE and ILE43VAL, while ZINC000016020583 was predicted to be not affected; furthermore, ZINC000016020583 had stronger binding tendency according to both docking and 10ns molecular dynamics simulation (Table 4).

## Discussion

We establish a hybrid system of classical and quantum computing to advance computer-aided drug design. It utilizes well-established virtual screening methods to identify lead compounds. It leverages machine learning to build an improved physical-statistical classifier capable of predicting the effect of variants from hypothetical and real mutations in receptor sequences of any genome to provide insights into drug efficacy. From screening to lead identification and to variant effect prediction, our approach can work not only in classical computing, but also in conjunction with quantum computing to eventually go beyond the current limits of classical computers. We demonstrated its performance and capability by applying the system to viral genomes with quantum machine learning (QML).

Antiviral drugs targeting SARS-CoV-2 3CL^pro^ could help to fight against the COVID-19 pandemic. As a proof of concept, we validated our screening method with the HCoV-229E 3CL^pro^ protein. We then applied our system to the SARS-CoV-2 3CL^pro^ protein, where two lead compounds, ZINC000016020583 and ZINC000036707984, were identified as potential inhibitors from ∼30,000 compounds subsampled from a drug dataset of over 11 million compounds. The new coronavirus is known to mutate frequently, sometimes with serious consequences in terms of immune evasion. It is essential to efficiently and reliably predict the effect of the mutations on any target drugs in the design process and to know the corresponding efficacy of a candidate drug. We therefore further applied our improved method of variant effect prediction to the identified compounds by using machine learning and both classical and quantum computing techniques, demonstrating that the performance of quantum computing is on par with classical computing. Our results showed indicative and insightful results based on ∼100 known mutations thus far, proving the utility of our approach for an advanced, promising, and robust drug design.

With the rapid advancement of computer power and algorithms, machine learning has become an indispensable tool to discover patterns in large-scale, high-dimensional data. We have explored the emerging field of quantum machine learning, which is the interplay of quantum computers and machine learning. Because of its exploitation of operations in the 2^N^-dimensional Hilbert space (for N qubits), quantum computing can solve certain problems that are hard for classical computers (https://quantumalgorithmzoo.org/; Shor, 1997; Van Dam et. al., 2006). In this work, we demonstrate several QML formulations trained on data that is highly imbalanced and is of lower quality than the test data, and then benchmark their performance on experimental data. The limited, orthogonal Platinum test data shows that the quantum ML models are at least on par with the established classical ML methods trained on the same dataset. Lastly, we assessed the fidelity of quantum simulators with respect to real quantum computers. Our classical-quantum hybrid approach has demonstrated how QC can be combined with classical computing robustly, with significant potential for added value in complex tasks such as protein-ligand binding predictions. As quantum computers become more powerful and stable, and more quantum-specific algorithms become available, we foresee that the utility of our approach will grow. For example, quantum computing can be used for protein design/folding (Perdomo et. al., 2008; Perdomo-Ortiz et. al. 2012), improving QM/MM calculation in molecular docking (Arodola et. al., 2017), modeling with neural-network quantum states (Choo et. al., 2020), as well as many other essential computational tasks.

