## Supplementary Materials for "Insights from Incorporating Quantum Computing into Drug Design Workflows"

### Supplementary Tables

| cQNN 3layer sim | | cQNN 1layer real | | cQNN 2layer real | | SNV identification | | | |
|---|---|---|---|---|---|---|---|---|---|
| prediction | exp val | prediction | exp val | prediction | exp val | position | ref | alt | lig |
| 0 | 0.007 | 0 | 0.328 | 0 | 0.407 | 160 | CYS | PHE | X77 |
| 1 | 0.998 | 1 | 0.534 | 1 | 0.531 | 168 | PRO | SER | X77 |
| 1 | 1 | 1 | 0.76 | 1 | 0.653 | 188 | ARG | SER | X77 |
| 0 | 0.003 | 0 | 0.262 | 0 | 0.468 | 160 | CYS | PHE | ZINC000016020583 |
| 1 | 0.998 | 1 | 0.56 | 1 | 0.63 | 168 | PRO | SER | ZINC000016020583 |
| 1 | 0.999 | 1 | 0.65 | 1 | 0.662 | 188 | ARG | SER | ZINC000016020583 |
| 0 | 0.001 | 0 | 0.168 | 0 | 0.384 | 160 | CYS | PHE | ZINC000036707984 |
| 1 | 0.999 | 1 | 0.723 | 1 | 0.579 | 168 | PRO | SER | ZINC000036707984 |
| 1 | 1 | 1 | 0.626 | 1 | 0.673 | 188 | ARG | SER | ZINC000036707984 |

**Table S1**. Results for the 3CL$^{pro}$ mutations run through the cQNN from Table 7, with expectation values (“exp val”) obtained from prediction (0 or 1 representing mutation impact) runs on IBM Quantum’s 5-qubit systems. The results shown include: a 3-layer (Farhi and Neven architecture) circuit where both training and predictions were run using the QASM simulator; and 1- and 2-layer circuits trained on the QASM simulator, but with prediction runs on an IBM Quantum 5-qubit system (marked by the term “real”). Orange cells are low-confidence predictions (with expectation values close to the threshold of 0.5).

| | | lig | X77 | | | | | ZINC000016020583 | | | | | ZINC000036707984 | | | | |
|---|---|---|---|---|---|---|---|---|---|---|---|---|---|---|---|---|---|
| | | Method | Genodock | Genodock w/ SVM | Weighted mQNN: 6-layer 0.15 margin | cQNN | qlsQNN | Genodock | Genodock w/ SVM | Weighted mQNN: 6-layer 0.15 margin | cQNN | qlsQNN | Genodock | Genodock w/ SVM | Weighted mQNN: 6-layer 0.15 margin | cQNN | qlsQNN |
| position | ref | alt | | | | | | | | | | | | | | | |
| 6 | MET | LEU | False | False | True | False | False | False | False | True | False | False | False | False | False | False | False |
| 8 | PHE | LEU | False | False | True | False | False | False | False | True | False | False | False | False | False | False | False |
| 15 | GLY | SER | False | False | True | False | False | False | False | True | False | False | False | False | False | False | False |
| 17 | MET | ILE | False | False | False | False | False | False | False | True | False | False | False | False | False | False | False |
| | | THR | False | False | False | False | False | False | False | True | False | False | False | False | False | False | False |
| 18 | VAL | ALA | False | False | False | False | False | False | False | True | False | False | False | False | False | False | False |
| 20 | VAL | ILE | True | True | True | True | True | False | False | True | False | False | False | False | False | False | False |
| 21 | THR | ILE | False | False | False | False | False | False | False | False | False | False | False | False | False | False | False |
| 23 | GLY | SER | False | False | False | False | False | False | False | True | False | False | False | False | False | False | False |
| 43 | ILE | VAL | True | True | True | True | True | True | False | True | False | False | True | True | True | True | True |
| 45 | THR | ILE | True | True | True | True | True | True | True | True | True | True | True | True | True | True | True |
| 46 | SER | PHE | True | True | True | True | True | True | True | True | True | True | True | True | True | True | True |
| | | PRO | True | True | True | True | True | True | True | True | True | True | True | True | True | True | True |
| 47 | GLU | LYS | False | False | False | False | False | False | False | False | False | False | False | False | False | False | False |
| 50 | LEU | PHE | True | True | True | True | True | True | True | True | True | True | True | True | True | True | True |
| 60 | ARG | CYS | False | False | False | False | False | False | False | False | False | False | False | False | False | False | False |
| 61 | LYS | ARG | False | False | True | False | False | False | False | True | False | False | False | False | False | False | False |
| 70 | ALA | SER | False | False | True | False | False | False | False | True | False | False | False | False | False | False | False |
| 71 | GLY | SER | False | False | True | False | False | False | False | True | False | False | False | False | False | False | False |
| 75 | LEU | PHE | False | False | True | False | False | False | False | True | False | False | False | False | True | False | False |
| 76 | ARG | SER | False | False | False | False | False | False | False | False | False | False | False | False | False | False | False |
| 78 | ILE | VAL | False | False | True | False | False | False | False | True | False | False | False | False | False | False | False |
| 80 | HIS | ASN | False | False | True | False | False | False | False | True | False | False | False | False | False | False | False |
| 81 | SER | PHE | False | False | False | False | False | False | False | False | False | False | False | False | False | False | False |
| 82 | MET | ILE | False | False | False | False | False | False | False | True | False | False | False | False | False | False | False |
| 83 | GLN | LYS | False | False | False | False | False | False | False | True | False | False | False | False | False | False | False |
| 88 | LYS | ARG | False | False | True | False | False | False | False | True | False | False | False | False | False | False | False |
| 89 | LEU | PHE | False | False | True | False | False | False | False | True | False | False | False | False | False | False | False |
| 90 | LYS | ARG | False | False | True | False | False | False | False | True | False | False | False | False | True | False | False |
| | | ASN | False | False | True | False | False | False | False | True | False | False | False | False | False | False | False |
| 94 | ALA | VAL | False | False | True | False | False | False | False | False | False | False | False | False | False | False | False |
| 96 | PRO | LEU | False | False | False | False | False | False | False | False | False | False | False | False | False | False | False |
| | | SER | False | False | False | False | False | False | False | True | False | False | False | False | False | False | False |
| 108 | PRO | LEU | False | False | False | False | False | False | False | False | False | False | False | False | False | False | False |
| | | SER | False | False | False | False | False | False | False | True | False | False | False | False | False | False | False |
| 116 | ALA | VAL | False | False | False | False | False | False | False | False | False | False | False | False | False | False | False |
| 121 | SER | LEU | False | False | False | False | False | False | False | False | False | False | False | False | False | False | False |
| 125 | VAL | ILE | False | False | True | False | False | False | False | True | False | False | False | False | False | False | False |
| 129 | ALA | VAL | False | False | False | False | False | False | False | False | False | False | False | False | False | False | False |
| 132 | PRO | LEU | False | False | False | False | False | False | False | False | False | False | False | False | False | False | False |
| | | SER | False | False | False | False | False | False | False | True | False | False | False | False | False | False | False |
| 135 | THR | ILE | False | False | False | False | False | False | False | False | False | False | False | False | False | False | False |
| 143 | GLY | SER | True | True | True | True | True | True | True | True | True | True | True | True | True | True | True |
| 157 | VAL | LEU | False | False | True | False | False | False | False | True | False | False | False | False | True | False | False |
| 160 | CYS | PHE | False | False | True | False | False | False | False | True | False | False | False | False | False | False | False |
| 162 | MET | ILE | False | False | False | False | False | False | False | False | False | False | False | False | False | False | False |
| 168 | PRO | SER | True | True | True | True | True | True | True | True | True | True | True | True | True | True | True |
| 178 | GLU | ASP | False | False | True | False | False | False | False | True | False | False | False | False | False | False | False |
| 184 | PRO | LEU | False | False | False | False | False | False | False | False | False | False | False | False | False | False | False |
| | | SER | False | False | False | False | False | False | False | False | False | False | False | False | False | False | False |
| 188 | ARG | LYS | True | True | True | True | True | True | True | True | True | True | True | True | True | True | True |
| | | SER | True | True | True | True | True | True | True | True | True | True | True | True | True | True | True |
| 190 | THR | ILE | True | True | True | True | True | True | True | True | True | True | True | True | True | True | True |
| 191 | ALA | VAL | True | True | True | True | True | True | True | True | True | True | True | True | True | True | True |
| 193 | ALA | SER | False | False | False | False | False | False | False | False | False | False | True | True | True | True | True |
| | | THR | False | False | False | False | False | False | False | False | False | False | True | True | True | True | True |
| | | VAL | False | False | False | False | False | False | False | False | False | False | True | True | True | True | True |
| 196 | THR | MET | False | False | True | False | False | False | False | True | False | False | False | False | False | False | False |
| 198 | THR | ILE | False | False | False | False | False | False | False | False | False | False | False | False | False | False | False |
| 199 | THR | ILE | False | False | False | False | False | False | False | False | False | False | False | False | False | False | False |
| 205 | LEU | VAL | False | False | True | False | False | False | False | True | False | False | False | False | False | False | False |
| 213 | ILE | VAL | False | False | False | False | False | False | False | False | False | False | False | False | False | False | False |
| 220 | LEU | PHE | False | False | False | False | False | False | False | False | False | False | False | False | False | False | False |
| 224 | THR | ALA | False | False | False | False | False | False | False | False | False | False | False | False | False | False | False |
| 225 | THR | ILE | False | False | False | False | False | False | False | False | False | False | False | False | False | False | False |
| 227 | LEU | PHE | False | False | True | False | False | False | False | False | False | False | False | False | False | False | False |
| 228 | ASN | SER | False | False | False | False | False | False | False | False | False | False | False | False | False | False | False |
| 232 | LEU | PHE | False | False | True | False | False | False | False | True | False | False | False | False | False | False | False |
| 234 | ALA | THR | False | False | True | False | False | False | True | True | False | False | False | False | False | False | False |
| | | VAL | False | False | False | False | False | False | False | False | False | False | False | False | False | False | False |
| 235 | MET | ILE | False | False | False | False | False | False | False | False | False | False | False | False | False | False | False |
| 236 | LYS | ARG | False | False | True | False | False | False | False | True | False | False | False | False | False | False | False |
| 241 | PRO | LEU | False | False | False | False | False | False | False | False | False | False | False | False | False | False | False |
| 244 | GLN | ARG | False | False | True | False | False | False | False | True | False | False | False | False | False | False | False |
| 246 | HIS | TYR | False | False | True | False | False | False | False | True | False | False | False | False | False | False | False |
| 247 | VAL | ILE | False | False | True | False | False | False | False | True | False | False | False | False | False | False | False |
| | | PHE | False | False | True | False | False | False | False | True | False | False | False | False | False | False | False |
| 249 | ILE | THR | False | False | False | False | False | False | False | False | False | False | False | False | False | False | False |
| 254 | SER | PHE | False | False | False | False | False | False | False | False | False | False | False | False | False | False | False |
| 255 | ALA | VAL | False | False | False | False | False | False | False | False | False | False | False | False | False | False | False |
| 257 | THR | ILE | False | False | False | False | False | False | False | False | False | False | False | False | False | False | False |
| 259 | ILE | THR | False | False | False | False | False | False | False | False | False | False | False | False | False | False | False |
| 260 | ALA | VAL | False | False | False | False | False | False | False | False | False | False | False | False | False | False | False |
| 261 | VAL | ILE | False | False | True | False | False | False | False | False | False | False | False | False | False | False | False |
| 263 | ASP | GLY | False | False | False | False | False | False | False | False | False | False | False | False | False | False | False |
| 264 | MET | VAL | False | False | False | False | False | False | False | False | False | False | False | False | False | False | False |
| 266 | ALA | VAL | False | False | False | False | False | False | False | False | False | False | False | False | False | False | False |
| 267 | SER | LEU | False | False | False | False | False | False | False | False | False | False | False | False | False | False | False |
| 274 | ASN | ASP | False | False | False | False | False | False | False | False | False | False | False | False | False | False | False |
| | | SER | False | False | False | False | False | False | False | False | False | False | False | False | False | False | False |
| 276 | MET | ILE | False | False | False | False | False | False | False | False | False | False | False | False | False | False | False |
| | | THR | False | False | False | False | False | False | False | False | False | False | False | False | False | False | False |
| 279 | ARG | CYS | False | False | False | False | False | False | False | False | False | False | False | False | False | False | False |
| 283 | GLY | ASP | False | False | True | False | False | False | False | True | False | False | False | False | False | False | False |
| 285 | ALA | VAL | False | False | False | False | False | False | False | False | False | False | False | False | False | False | False |
| 298 | ARG | LYS | False | False | True | False | False | False | False | True | False | False | False | False | False | False | False |
| 301 | SER | LEU | False | False | False | False | False | False | False | False | False | False | False | False | False | False | False |
| 304 | THR | ILE | False | False | False | False | False | False | False | False | False | False | False | False | False | False | False |

**Table S2**. Impact of 6W63 mutations on the protein ligand binding as predicted by different methods. True means such mutation would negatively affect the binding.

| Truth | Genodock | Genodock w/ SVM | cQNN | qisQNN | Weighted mQNN: 3-layer 0.15 margin | Weighted mQNN: 3-layer 0.5 margin | Weighted mQNN: 6-layer 0.15 margin | Weighted mQNN: 6-layer 0.5 margin |
|---|---|---|---|---|---|---|---|---|
| 1 | 0.93 | 0.9268569507198795 | 0.99 | 1.0 | 0.1830009520053863 | 0.7894039154052734 | 0.2162638008594513 | 0.7903943657875061 |
| 1 | 0.98 | 0.9555163944734489 | 1.0 | 1.0 | 0.2643681764602661 | 0.982441246509552 | 0.287376120686531 | 1.019123762845993 |
| 1 | 0.95 | 0.9488339945746942 | 1.0 | 1.0 | 0.2680611312389374 | 1.045918583869934 | 0.2830354422330856 | 1.0903245210647583 |
| 1 | 0.94 | 0.9469362136735343 | 1.0 | 0.99 | 0.1604686081409454 | 0.7895245850086212 | 0.1817051023244857 | 0.8037940263748169 |
| 1 | 0.86 | 0.9425919635153927 | 1.0 | 1.0 | 0.1607889533042907 | 0.8416368961334229 | 0.1862980499863624 | 0.8347724676132202 |
| 1 | 0.9 | 0.9236602491670737 | 1.0 | 1.0 | 0.1840356588363647 | 0.8333681225776672 | 0.2194388210773468 | 0.8431910574436188 |
| 1 | 0.36 | 0.12225507338418928 | 0.01 | 0.0 | -0.2295016050338745 | -0.4152374267578125 | -0.3419502945616841 | -0.7684855461120605 |
| 1 | 0.82 | 0.9328003382255063 | 1.0 | 1.0 | 0.1337878406047821 | 0.9259080290794371 | 0.212616354227066 | 0.932088017463684 |
| 1 | 0.94 | 0.9326224619811532 | 1.0 | 1.0 | 0.3136352598667145 | 0.886750340461731 | 0.3688510730862617 | 0.933920294046402 |
| 1 | 0.46 | 0.04336523345411437 | 0.0 | 0.0 | -0.2710021436214447 | -0.5820097476243973 | -0.3925638049840927 | -0.8861829191446304 |
| 1 | 0.89 | 0.9257558728536626 | 0.98 | 1.0 | 0.3519052267074585 | 0.9717921018600464 | 0.4044060409069061 | 0.9879558980464936 |
| 0 | 0.98 | 0.9291943609725529 | 1.0 | 1.0 | 0.3034542500972748 | 1.001962125301361 | 0.3247562535107136 | 1.065472811460495 |
| 1 | 0.98 | 0.9294476857297507 | 1.0 | 1.0 | 0.3121184110641479 | 1.0087187886238098 | 0.3372189179062843 | 1.0769792199134829 |
| 1 | 0.98 | 0.9293800099002388 | 1.0 | 0.99 | 0.312018558382988 | 1.008864939212799 | 0.3370962888002395 | 1.0769537687301636 |
| 0 | 0.98 | 0.9036113390180823 | 0.99 | 1.0 | 0.1210971176624298 | 0.5470504462718964 | 0.1576818823814392 | 0.5360137075185776 |
| 1 | 0.98 | 0.9322069018429289 | 0.99 | 1.0 | 0.2501252219080925 | 0.8325046300888062 | 0.3370108306407928 | 0.9296062290668488 |
| 1 | 0.84 | 0.925366777314634 | 0.99 | 1.0 | 0.17581707239151 | 0.6961102187633514 | 0.2483108788728714 | 0.7734333872795105 |
| 1 | 0.07 | 0.1039628834092077 | 0.02 | 0.0 | -0.3808790594339371 | -0.4994791895151138 | -0.3537298366427421 | -0.7296166121959686 |
| 0 | 0.02 | 0.0576159802424963... | 0.01 | 0.0 | -0.2032391279935836 | -0.5760806798934937 | -0.1227966845035553 | -0.5578334778547287 |
| 0 | 0.12 | 0.18169240870196565 | 0.0 | 0.0 | -0.1952921599149704 | -0.5437014400959015 | -0.1578945070505142 | -0.5857659243047237 |
| 0 | 0.91 | 0.9355384307328594 | 0.99 | 1.0 | 0.5089084059000015 | 1.3761892914772034 | 0.602888286113739 | 1.5492733716964722 |
| 1 | 0.79 | 0.3640424881984206 | 0.02 | 0.0 | -0.2664129137992859 | -0.8431113883852959 | -0.271107017993927 | -0.5070528835058212 |
| 0 | 0.68 | 0.34858353185419577 | 0.03 | 0.0 | -0.2816145122051239 | -0.8880771696567535 | -0.3094122558832168 | -0.6160365343093872 |
| 0 | 0.38 | 0.20185649158200228 | 0.0 | 0.01 | -0.0572662949562072 | -0.2601195201277733 | 0.0317667722702026 | -0.3045706748962402 |
| 1 | 0.99 | 0.9271782112166932 | 1.0 | 1.0 | 0.2744050323963165 | 0.8843718767166138 | 0.299609512090683 | 0.9518534541130066 |
| 1 | 0.96 | 0.9279040726767133 | 1.0 | 1.0 | 0.2840185016393661 | 0.8922819495201111 | 0.3127017915248871 | 0.965608298778534 |
| 1 | 0.97 | 0.9281234346980679 | 1.0 | 1.0 | 0.2843588888645172 | 0.8921630382537842 | 0.3130905330181122 | 0.965913623571396 |
| 1 | 0.97 | 0.9281234346980679 | 1.0 | 1.0 | 0.2843588888645172 | 0.8921630382537842 | 0.3130905330181122 | 0.965913623571396 |
| 1 | 1.0 | 0.9354164686314875 | 1.0 | 1.0 | 0.2361021190881729 | 0.7494884133338928 | 0.2461705952882766 | 0.7750895321369171 |
| 0 | 0.86 | 0.9200141642810918 | 0.99 | 1.0 | 0.0618825256824493 | 0.5603171288967133 | 0.1954734176397323 | 0.6490544378757477 |
| 0 | 0.97 | 0.9514060296739066 | 1.0 | 1.0 | 0.3297361433506012 | 1.118439257144928 | 0.4296564236283302 | 1.2498580813407898 |
| 0 | 0.88 | 0.9422936728694619 | 0.99 | 1.0 | 0.2696277946233749 | 1.0044042468070984 | 0.3441464006900787 | 1.085256963968277 |
| 1 | 1.0 | 0.9542892239501612 | 1.0 | 1.0 | 0.3199249356985092 | 1.0453458428382874 | 0.4016116484999656 | 1.1605996787548063 |
| 1 | 0.81 | 0.9138550014542158 | 0.99 | 1.0 | 0.2645938992500305 | 0.7561655640602112 | 0.3093237578868866 | 0.860230565071106 |
| 1 | 0.13 | 0.0579900331910387... | 0.02 | 0.0 | -0.3215516600757837 | -0.7039915770292282 | -0.4492231048643589 | -0.9280913472175598 |
| 0 | 0.99 | 0.9474381453417254 | 0.99 | 1.0 | 0.2964816614985466 | 0.9193879961967468 | 0.3163446933031082 | 0.979468047618866 |
| 1 | 0.95 | 0.9498720635982024 | 0.99 | 1.0 | 0.2122353017330169 | 0.4541453421115875 | 0.3114438652992248 | 0.5849291384220123 |
| 1 | 0.99 | 0.9403861358779589 | 1.0 | 1.0 | 0.1874006241559982 | 0.5855906009674072 | 0.2080089896917343 | 0.6528389751911163 |
| 1 | 0.94 | 0.9009064419117118 | 1.0 | 1.0 | 0.1797963045537471 | 0.4416861832141876 | 0.2084318213164806 | 0.470931202173233 |
| 1 | 0.96 | 0.903087803409521 | 1.0 | 1.0 | 0.1821930408477783 | 0.4452338218688965 | 0.2161457054316997 | 0.4732409715652466 |
| 1 | 1.0 | 0.9304028098026755 | 0.99 | 1.0 | 0.1616241931915283 | 0.5514388382434845 | 0.2123548090457916 | 0.5784996449947357 |
| 1 | 0.9 | 0.9207445970427767 | 0.99 | 1.0 | 0.2762935757637024 | 0.6961502730846405 | 0.2453422397375106 | 0.6800020039081573 |
| 1 | 0.54 | 0.10666518652586217 | 0.0 | 0.01 | -0.4996291846036911 | -0.6235489249229431 | -0.2904951870441437 | -0.8405472934246063 |
| 1 | 0.86 | 0.8741195527169839 | 1.0 | 1.0 | -0.1139766424894332 | 0.2508638501167297 | 0.0105920135974884 | 0.2914774119853973 |
| 0 | 0.38 | 0.12046748496905657 | 0.0 | 0.0 | -0.4348517507314682 | -0.5029818415641785 | -0.226512610912323 | -0.6941833794116974 |
| 1 | 0.52 | 0.12396074317614929 | 0.01 | 0.0 | -0.4583893269300461 | -0.5857871770858765 | -0.2528710663318634 | -0.7703317403793335 |
| 1 | 0.85 | 0.851838192574697 | 0.99 | 1.0 | -0.129429891705513 | 0.1921952366828918 | -0.0007209777832031 | 0.2478602528572082 |
| 1 | 0.9 | 0.8944499365649036 | 0.99 | 1.0 | -0.1062480583786964 | 0.2872803248465061 | 0.0364325940608978 | 0.3413818925619125 |
| 1 | 0.98 | 0.956704044595553 | 0.99 | 1.0 | 0.2767821997404098 | 0.926838517189026 | 0.2987268716096878 | 0.9671456515789032 |
| 0 | 0.95 | 0.9497711825347309 | 1.0 | 1.0 | 0.2903060764074325 | 0.987718105316162 | 0.3031914234161377 | 1.033528059720993 |
| 1 | 0.97 | 0.9483943370487965 | 1.0 | 1.0 | 0.3092516660690307 | 1.0773200988769531 | 0.3367226719856262 | 1.1099363267421722 |
| 1 | 0.95 | 0.9479232965540983 | 0.99 | 1.0 | 0.1842129975557327 | 0.7370850145816803 | 0.1964149549603462 | 0.7582097053527832 |
| 1 | 0.95 | 0.9040954177617331 | 0.99 | 1.0 | 0.185416642576456 | 0.4490920901298523 | 0.2162258718162775 | 0.4774225652217865 |
| 1 | 0.94 | 0.9040057345607385 | 1.0 | 1.0 | 0.1821879893541336 | 0.449773371219635 | 0.2198563069105148 | 0.4633449912071228 |
| 1 | 0.98 | 0.9334732568455961 | 0.99 | 0.99 | 0.1604543626308441 | 0.5346464812755585 | 0.2123931348323822 | 0.5615296214818954 |
| 1 | 0.95 | 0.9400336969747339 | 1.0 | 1.0 | 0.1250978112220764 | 0.5127823650836945 | 0.2394162714481353 | 0.6154802143573761 |
| 1 | 0.95 | 0.9400336969747339 | 1.0 | 1.0 | 0.1250978112220764 | 0.5127823650836945 | 0.2394162714481353 | 0.6154802143573761 |
| 1 | 0.88 | 0.841645547871589 | 1.0 | 1.0 | 0.031851440668106 | 0.1402499079704284 | 0.0579235553741455 | 0.1578415036201477 |
| 1 | 0.88 | 0.841645547871589 | 1.0 | 1.0 | 0.031851440668106 | 0.1402499079704284 | 0.0579235553741455 | 0.1578415036201477 |
| 1 | 0.93 | 0.9033593348558416 | 1.0 | 1.0 | 0.0681804418563842 | 0.313897609710693 | 0.1637658774852752 | 0.381616935133934 |
| 1 | 0.93 | 0.9033593348558416 | 1.0 | 1.0 | 0.0681804418563842 | 0.313897609710693 | 0.1637658774852752 | 0.381616935133934 |
| 1 | 0.96 | 0.9028868131193819 | 1.0 | 1.0 | 0.0835889279842376 | 0.339255541563034 | 0.1478975713253021 | 0.3938570320606231 |
| 1 | 0.96 | 0.9028868131193819 | 1.0 | 1.0 | 0.0835889279842376 | 0.339255541563034 | 0.1478975713253021 | 0.3938570320606231 |
| 1 | 0.87 | 0.9366365756342846 | 1.0 | 1.0 | 0.5314924567937851 | 1.3708425760269165 | 0.6029666066169739 | 1.479819655418396 |
| 1 | 0.9 | 0.9357978635889364 | 1.0 | 1.0 | 0.2169839739799499 | 0.5046365559101105 | 0.2345022708177566 | 0.4796957820653915 |
| 1 | 0.11 | 0.14356709214722393 | 0.01 | 0.02 | -0.2565460652112961 | -0.5512207746505737 | -0.1943202018737793 | -0.6515644192695618 |
| 1 | 0.25 | 0.1583763387361935 | 0.0 | 0.0 | -0.2675242722034454 | -0.5801892280578613 | -0.2366800010204315 | -0.6748873293399811 |
| 1 | 0.16 | 0.14765055434662772 | 0.01 | 0.0 | -0.2812144458293915 | -0.584501102566719 | -0.2670266628265381 | -0.7010801434516907 |
| 0 | 0.3 | 0.18236426768666814 | 0.0 | 0.0 | -0.2716587036848068 | -0.5802980363368988 | -0.2510666102170944 | -0.6824607849121094 |
| 1 | 0.94 | 0.90335397606952 | 1.0 | 1.0 | 0.182493295520544 | 0.4489427506923675 | 0.215595512650907 | 0.4677069783210754 |
| 1 | 0.96 | 0.9056172271092526 | 1.0 | 1.0 | 0.1873328536748886 | 0.453334629535675 | 0.2244866788387298 | 0.4744450747966766 |
| 1 | 0.99 | 0.933645231279453 | 1.0 | 1.0 | 0.1587545871734619 | 0.5277284383773804 | 0.2098719328641891 | 0.5528465360403061 |
| 1 | 0.98 | 0.925810695214788 | 1.0 | 1.0 | 0.2712032496929168 | 0.718929648399353 | 0.3040393963456154 | 0.7738063037395477 |
| 1 | 0.98 | 0.927351597156782 | 0.99 | 1.0 | 0.2716707289218902 | 0.7049286961555481 | 0.3051925525069237 | 0.7558083832263947 |
| 1 | 0.98 | 0.9301710928098433 | 0.98 | 1.0 | 0.2873266637325287 | 0.8419066071510315 | 0.3356474414467811 | 0.9273084700107574 |
| 0 | 0.98 | 0.9802431763232407 | 1.0 | 1.0 | 0.3180528879165649 | 0.748442530632019 | 0.2335161715745926 | 0.8032137155532837 |
| 0 | 0.96 | 0.9726223160865118 | 0.98 | 1.0 | 0.2706646919250488 | 0.6030084192752838 | 0.1953307017683982 | 0.6322290599346161 |
| 1 | 0.96 | 0.9435485568815717 | 0.01 | 0.0 | 0.1703386902809143 | 0.2690293192863464 | 0.1793813407421112 | 0.4637284576892853 |
| 0 | 0.63 | 0.07637452873776687 | 0.0 | 0.0 | -0.234207272529602 | -0.4862769842147827 | -0.3704657107591629 | -0.6367243528366089 |
| 0 | 0.71 | 0.07309213330516161 | 0.01 | 0.0 | -0.2803738415241241 | -0.5263530761003494 | -0.5128955617547035 | -0.658849872648716 |
| 0 | 0.52 | 0.05056800686538992 | 0.02 | 0.0 | -0.4028680026531219 | -0.6821742355823517 | -0.8772078156471252 | -1.1883766651153564 |
| 0 | 0.7 | 0.0791113337305376 | 0.01 | 0.0 | -0.275914192199707 | -0.5183612108230591 | -0.4130439460277557 | -0.605637788772583 |
| 1 | 0.98 | 0.9472312520323712 | 0.98 | 1.0 | 0.4379953444004059 | 1.1257745623588562 | 0.4806888997554779 | 1.2501358389854431 |
| 1 | 0.95 | 0.9277508967008009 | 1.0 | 1.0 | 0.2733129411935806 | 0.8326157927513123 | 0.2957621663808822 | 0.8453674912452698 |
| 1 | 0.44 | 0.06906736483036749 | 0.0 | 0.01 | -0.2074703276157379 | -0.5430327281355858 | -0.3122403472661972 | -0.7505329698324203 |
| 1 | 0.52 | 0.0601982723696853... | 0.0 | 0.01 | -0.3715180456638336 | -0.6635741963982582 | -0.570472739636898 | -1.0179263949394226 |

**Table S3**. Prediction scores as output by various methods. The Genodock scores are recorded from the predict_proba function in Python's Scikit-Learn package. The scores of quantum methods are computed from repeated measurements of the z-projection of the output qubit. We note that the projected measurement is not strictly linear in terms of probability due to its nature of being a z-projection of a "spin" that can be rotated about the X/Y axis.

### Supplementary Methods

#### GenoDock data and feature groups

The construction of the GenoDock dataset used in our analyses is detailed in the original publication (Wang et. al., 2019) . We briefly summarize the procedure. The starting point was to find proteins with experimental co-crystal structures that included at least one FDA-approved drug ligand from a list of 113 drugs. The net result was a set of 228 proteins. Non-synonymous germline single-nucleotide variants (SNVs) from the Exome Aggregation Consortium (Lek et. al., 2016) and somatic SNVs from The Cancer Genome Atlas (Cancer Genome Atlas Research Network, 2012; Cancer Genome Atlas Research Network, 2013; Cancer Genome Atlas Research Network, 2016) were mapped on to the 228 proteins, using a previously published pipeline (Kumar, 2016). The set of successful mappings led to 8,565 ExAC SNVs and 1,718 TCGA SNVs. The resulting database thus includes 10,283 non-synonymous human SNVs across 228 proteins. Each of these SNVs was associated with a single amino acid residue within the corresponding PDB structures for the target proteins, and the "wild-type" PDB structure of the protein was mutated at this site, using the program Modeller (Webb and Sali 2016) (resulting in a "mutant" structure). To generate the so-called "pseudo gold standard" binding affinities for the SNV-Protein structure-Ligand sets, the program AutoDock Vina (Trott, 2010) was used to rigidly dock the ligand to both the wild-type and mutant structures, and calculate the resulting binding affinity scores. The binding affinity of interest was the difference in binding affinities between the mutant and wild-type dockings. To further remove any biases associated with the docking program, only the sign of the binding affinity difference was considered in the downstream training, where a positive change in binding affinity was considered "disruptive" while a negative or zero magnitude change in binding affinity was considered to be a "non-disruptive" or "neutral" change. The original GenoDock model was a RandomForest trained on this dataset.

The feature groups considered in the original paper were divided according to the particular component of the SNV-Protein structure-Ligand triplet they are associated with:

1. **SNV features:** allele frequency; SIFT (Kumar, 2009); PolyPhen-2 (Adzhubei, 2013); and GERP scores (Davydov et. al. 2010)
2. **Ligand Features:** Molecular weight; H-bond donor and acceptor counts (two separate features); rotatable bond count; polar surface area.
3. **Structure-associated features:** Amino acid side-chain volume change index (log base-2 of the ratio of the mutant amino acid volume to the wild-type amino acid volume); distance between the mutation and drug ligand (shortest distance of the mutated residue to a heavy atom of the ligand); polarity change index (the mutation's change of residue polarity index, which is +/-1 for charged residue, 0.5 for polar, and 0 for hydrophobic amino acid); binding site "on/off" (if the distance as defined previously is less than 8 Angstroms = "on"; else = "off").

These feature groups were then used in 4 different combinations to reflect variation in the amount of information available to a user:

I. **Feature group 1:** SNV features only (4 features in total)
II. **Feature group 2:** SNV + Structure features - Distance feature (7 features in total)
III. **Feature group 3:** SNV + Ligand features + Binding site “on/off” feature (10 features in total)
IV. **Feature group 4:** SNV + Structure + Ligand features (13 features in total)

These feature groups are also employed in our current analysis to enable one-to-one comparisons with the original GenoDock model.

While the paper focused on human proteins, we assessed the suitability of some genomic features for viral genomes. Specifically, we considered features such as SIFT (Kumar, 2009), PPH (Adzhubei, 2013), and GERP (Davydov et. al. 2010) scores, and germline/somatic allele frequency. The variant annotation of well-studied and -tracked viruses such as SARS-CoV-2 is feasible (see variant frequencies and SIFT scores in Dunham et. al. (2021)). However, we removed such features to build non-human-genome-compatible classifiers applicable to a wider variety of viruses lacking sufficient information. Furthermore, feature space reduction also facilitates downstream calculations on the limited-qubit quantum computers considered. In general, though, the methods are designed to incorporate all features given availability of sufficiently comprehensive variant databases.

In the current study, we specifically investigate nonhuman features in the interest of applying the QNN framework to the SARS-CoV-2 $3CL^{pro}$ protein. That is, we consider the subset of all features not specific to human genomes (namely everything except the GERP, PolyPhen2, and SIFT scores, and the allele frequency). This left us with nine features (*bind_site*, *distance*, *molecular_weight*, *H_bond_donor*, *H_bond_acceptor*, *rotatable_bond*, *Polar_Surface_Area*, *polarity_change_index*, and *volume_change_index*), which was a tenable number for simulated QNNs. For QNNs that run on IBM’s 5-qubit systems, we subselected again to those features which are not ligand-specific, leaving us with four features: *bind_site*, *distance*, *polarity_change_index*, and *volume_change_index*. This is a 5-qubit problem (one input qubit for each feature plus an additional readout qubit) and therefore perfectly suited to our resources. It is worth noting that, in our testing on simulated devices, we found no significant performance difference between the nine-feature group and the four-feature group. We suspect this is due to the dominance of the *bind_site* feature in the prediction process, but, regardless of the cause, it reassures us that working with the four-feature group is still a meaningful problem.

As explained below, the necessity of balancing the datasets (in terms of output labels) for the training procedure resulted in different dataset sizes for different feature groups. This is because removing some of the features resulted in multiple data points being redundant in the resulting lower-dimension space, and we removed such redundant data points. The partition sizes (i.e. the size of the equally distributed sets of data points for the training, validation and testing groups) are (1) Feature group 1: 150; (2) Feature group 2: 400; (3) Feature group 3: 360; (4) Feature group 4: 440; and (5) Viral feature group (excluding human-specific features): 440.

#### Platinum dataset

We also tested our QNN models on an external dataset to get an idea of its generalizability, using the same Platinum dataset (Pires et. al. 2015) as the original GenoDock paper. The Platinum datasets consist of experimental binding affinities for target proteins before and after mutations. The GenoDock paper retained only human protein and point mutations (i.e. single amino acid mutations) for downstream analysis. Further filtering steps include: removing mutations that do not arise from single-nucleotide variants and removing those mutations that could not be mapped onto their associated UniProtKB canonical amino acid sequences. The net result was a set of 86 unique mutated-protein + ligand pairs. A similar binarization as in the GenoDock dataset was carried out, where a mutation was disruptive if the change in binding affinity upon mutation was positive, and non-disruptive otherwise. After processing, the data set has 66 positives and 20 negative data points.

#### cQNN and qisQNN data preprocessing

##### 1. Balancing the data

After being appropriately binarized or rescaled, we then balanced our datasets in terms of label values. We accomplished this by first splitting the data according to label, then rebuilding the dataset one input at a time, alternating between each label category. This alternation process halted as soon as one label category was depleted, so the final dataset was always perfectly balanced. The data can be shuffled (randomly or according to a specific seed) and if shuffling is desired, it is executed before the alternation process. By doing so, we ensured that the label pattern was not disrupted.

##### 2. Partitioning into training/testing/validation sets

Finally, the data was split into partitions for training, validation, and testing. Because the dataset was already ordered such that the labels alternated, it was easy to keep the partitions balanced; as long as the partition size is an even number, it is guaranteed that each partition has a balanced number of inputs labeled “0” and “1.” We began with an even 1:1:1 split of training:validation:testing, and after comparison to other partition ratios found the even split to be the most effective at preventing overfitting and maintaining generalizability. Moreover, the size of each partition was chosen to be the maximum allowed by each balanced data subset. These choices are adjustable according to the partition ratio and partition size hyperparameters.

### Quantum Neural Networks

#### 1. Architecture

The layers of the circuit consist of the sequential application of a series of rotation gates. As mentioned in the main manuscript, we chose to focus on the application of 2-qubit gates to combinations of qubits. Specifically, we alternated the application of $R_{ZX}(\theta) = exp\left(i\theta\, Z_{input} \otimes X_{readout}\right)$ and $R_{XX}(\theta) = exp\left(i\theta\, X_{input} \otimes X_{readout}\right)$, where a single layer consisted of the application of such gates to each input in turn. These gates can be represented as matrices in the 4 X 4-dimensional 2-qubit space.

#### 2. Fitting and Classification

There are three stages to fitting the QNN models: training, validation, and final testing. This three-part process is meant to prevent the models from overfitting to the training data and to get a good idea of generalization to unseen data.

##### *2a. TRAINING*

For training, the QNN is fed all of the input data from the train partition of the dataset in batches, the average loss is computed for each batch, and the θ vector is updated accordingly. We use batch stochastic gradient descent optimization, with the batch size and gradient shift size being hyperparameters. We use a batch size of 20 and a shift size of pi/2 (see below).

Before the model can begin learning, all $R_{ZX}$ and $R_{XX}$ gates must be initialized with some angle $\theta_n$. We tested both random initializations where every parameter was unique and uniform initializations where all parameters from the same value and found that performance was often similar. That said, there were occasionally random initializations from which the model was unable to learn. Additionally, we thought it would be useful to explore how much each $\theta_n$ in fully trained models deviates from a shared initial value and to gain insight into interpretability or at least parameter significance. Thus, we decided to initialize all models from uniform parameters. To determine the best initial parameters, we looked at a selection of feature groups over a number of cases. For those feature groups, we divided the parameter space between 0 and pi as increments of pi/8 and initialized models by setting all parameters to the given value. We saw no significant difference in performance between the eight models regardless of their starting parameters. We ultimately decided to initialize all $\theta_n$ to a starting value of π/2, choosing this value because it creates maximum entanglement in the $R_{ZX}$ and $R_{XX}$ gates. This starting value can be adjusted with the initial parameters hyperparameter.

Parameter update is governed by the loss function (specifically, by the gradient of the loss function evaluated for the given θ vector). Farhi and Neven calculate the loss according to $loss(\vec{\theta}, z) = 1 - l(z)\langle z, 1\left|U^{\dagger}(\vec{\theta})Y_{n+1}U(\vec{\theta})\right|z, 1\rangle$ (where l(z) is the actual label value, which is multiplied by the prediction as defined above) and propose a method to calculate the gradient thereof analytically.

In our implementation, we use the same loss function but calculate the gradient via a finite difference approximation. $\frac{df}{dx}(x) = \frac{f(x+\epsilon) - f(x-\epsilon)}{2\epsilon} + O(\epsilon^2)$ This method is referred to by Farhi and Neven (Farhi et. al., 2018) and detailed in (Mitarai, 2019). Using this method, the gradient of the loss function is approximated by shifting each $\theta_n$ down by some value ϵ (which is dictated by shift size) and calculating the loss, shifting up by that value and calculating the loss again, then taking the difference between those values and dividing by the shift. (Crooks, 2019) and the IBM Q tutorials (https://qiskit.org/documentation/tutorials/operators/02_gradients_framework.html) suggest a shift size of pi/2, and our own explorations confirm this as the most effective. We tested shift sizes from 0 to pi/2 in intervals of 0.25, and none of the other shift values performed better than the original pi/2. The PyTorch modules we used to construct our QNN did not support quantum circuits with built-in functions, so the finite difference approximation was computed with an external function.

The gradient-calculation step is the most computationally demanding part of the QNN, for it involves calculating the expectation value of the circuit 2L*shots times, where L is the length of the θ vector and shots is the number of times the circuit is executed to approximate the expectation value. We offset this computational load by parallelizing the forward and backward pass of each data batch through the QNN. Depending on the number of CPUs allotted to training, several data inputs in each batch are passed through the QNN simultaneously. The loss and gradient thereof for each batch are computed independently on each thread. Once every input in the batch is seen, the threads are rejoined and the average loss and average gradient of the loss is calculated for the entire batch.

Once the gradient of the loss is calculated, all parameters in the θ vector are updated according to $\vec{\theta} \rightarrow \vec{\theta} - r\left(\frac{loss(\vec{\theta}, z)}{\vec{g}^2}\right)\vec{g}$ (where r is the learning rate hyperparameter and g is the gradient of the loss function), and the QNN progresses to the next batch of data. After a few epochs (the exact number is governed by decay start), the learning rate used in parameter update decays with each successive epoch as it is multiplied by some decay rate. Using independent grid searches for each hyperparameter, we find learning rate=0.1, decay start=5, and decay rate=0.9 to be the most effective for learning without overfitting.

##### *2b. VALIDATION*

After every epoch, the partially-trained QNN is validated against a previously unseen dataset. The partially-trained parameters are fixed for the entire validation step, and the model attempts to classify all of the validation partitions. The average loss across the partition is then calculated and saved alongside the corresponding model. Finally, the QNN returns to the training stage and the parameters are once again updated accordingly.

These two steps are repeated a number of times according to the epochs hyperparameter, and each time the validation loss is calculated, it is compared with the previously saved model. If the current model performs better than the saved model, it is cataloged instead. If not, the current model is fed the training partition again in the hopes of training it further. If, for several epochs in a row the current model cannot outperform the saved model, the fitting routine is exited and the saved model is presented as the best possible version of the QNN. The number of epochs attempted before exiting the training routine is determined by the patience hyperparameter.

##### *2c. TESTING*

Finally, this fully-trained QNN is evaluated on the test partition, which is another set of previously unseen data. As the QNN attempts to classify all of the test data, a number of metrics are calculated, including the test loss, precision, accuracy, and the area under the ROC curve (AUROC). Aside from the test loss (which is calculated using the loss function outlined above), all evaluation metrics are imported from scikit-learn (see EVALUATION).

#### 3. Evaluation

Because QNN classification is based on the expectation value of a quantum measurement, the predictions are stochastic. Evaluating models can therefore be a bit nuanced, as running the same model on the same test data multiple times might result in slightly different predictions. The previously mentioned test loss (the average of the loss function evaluated across the entire test partition) is a good starting point for evaluating models, but it can fluctuate arbitrarily and is best supplemented by other metrics as well. We have evaluated several such metrics and describe them in the following.

##### *3a. ROC AND AUROC*

One metric we have used for evaluating our models is the receiver operating characteristic (ROC) curve. Since the QNN’s label predictions are computed from an expectation value, they can be any value between 0 and 1. Those labels must then be binarized, but the best threshold for binarization is not obvious. Here lies the strength of the ROC curve: it compares the rates of false and true positive classifications across a range of thresholds, providing a dynamic understanding of a model’s performance. This performance can be quantified by calculating the area under the curve (AUC), where a value of 1 corresponds to perfect classification and a value of 0.5 is roughly equivalent to guessing. To plot the ROC curves and calculate the AUROC, we imported functions from scikit-learn.

##### *3b. CONFUSION MATRIX*

To supplement the ROC curves, we also calculated the confusion matrix for each model’s classification of the test partition (again using scikit-learn). This matrix describes the rates of true positives, false positives, true negatives, and false negatives across the dataset. Those rates can then be used to calculate other metrics such as precision and accuracy, which allow a more granular understanding of model performance. However, calculating the confusion matrix requires binary label values, so the QNN's continuous predictions must be thresholded at some value. Unlike with the ROC curve, we can't easily evaluate a broad range of thresholds, so we chose to use 0.5 for this binarization.

#### 4. Overall Model Implementation

##### *4a. HYPERPARAMETERS*

Because our QNN is largely custom-built, there are a multitude of hyperparameters to adjust and optimize. As previously mentioned, these include batch and partition size, the number of shots (i.e. times the circuit is executed each run to calculate the expectation value), shift size to be used in gradient approximation, and patience in searching for a best model. For each of these hyperparameters, we ran a grid search across possible values and selected those which led to best model performance. Specific values for all hyperparameters are listed in the RESULTS section and are set as defaults in the QNN source code. That said, all hyperparameters are still easily adjustable as options used when calling the QNN script.

##### *4b. QISKIT'S MACHINE LEARNING MODULE*

After the development of our custom QNN, Qiskit released a machine learning module which includes a NeuralNetworkClassifier class and automates much of the neural network construction process. Instead of constructing the network from the quantum circuit up, the user only needs to provide a feature map to encode data to the circuit and a circuit architecture for use in classification. There are a number of pros and cons to using this module. The most obvious advantage is ease of use: a NeuralNetworkClassifier automatically handles computations such as loss calculation and parameter update, and a wide variety of Qiskit functions make it trivial to test other parameter optimization methods. That said, this functionality comes at the expense of control, especially with regard to model validation. To make the Qiskit training conform to the same validation framework as we have used in our custom module, we had to build a wrapper around the NeuralNetworkClassifier. Ultimately, our additions to the validation process amount to rebuilding the neural network from more primitive classes, which we have already done in our custom implementation. For this reason, we have primarily used Qiskit's NeuralNetworkClassifier as a supplement to our own QNN, using it to explore optimization methods and determine which of those is worth adapting to our custom implementation.

In building QNN models with Qiskit's NeuralNetworkClassifier ("qisQNNs"), we chose to use their Constrained Optimization BY Linear Approximation (COBYLA) optimizer, implemented using the Qiskit default parameters. This optimization is not the exact same as the stochastic gradient descent used in our custom QNN ("cQNN") models, so when judging model performance, we'll take into consideration possible effects due to the differences in parameter updates. That said, we don't expect significant impacts on performance aside from training times. This holds true for data batching as well; whereas the cQNN is fed data in batches of 20, the qisQNN is given access to the entire training partition at once.

As the module is written, Qiskit's NeuralNetworkClassifier trains on a single data partition and then uses that trained model to make predictions on the test data. This compares poorly with the cQNN, which trains on one partition and then validates on another several times before ever seeing the test data. Indeed, evaluating qisQNN models as written saw them consistently performing worse than cQNN models, assumedly because they were seeing less data during training. To remedy this, we have incorporated a validation stage into the qisQNN training procedure.

Validation for the qisQNN mirrors that of the cQNN, but the routine is external to the training routine instead of happening alongside it. Before validation, the qisQNN is trained on the training partition according to Qiskit's protocol. Then, the trained model is scored against the validation partition, and that score is archived alongside the model parameters. The qisQNN is then warm-started to learn again on the training partition, and once finished, it is once again scored on the validation partition. The validation score is then compared to the archived score. If the new model performs better, it's archived instead as the new best model. If not, the training is repeated with the same model in the hopes of improving performance. Like in the cQNN validation, this process is repeated a number of times, and if the qisQNN's validation score does not improve for a number of iterations in a row, the routine is exited and the best saved model is returned as the fully-trained qisQNN. This validation routine has improved the qisQNN's performance drastically, elevating it to the level of the cQNN and sometimes even surpassing it. (see RESULTS).

##### *4c. CHOOSING BETWEEN SIMULATIONS AND RUNNING ON QUANTUM COMPUTERS*

The vast majority of our QNN models were trained and executed using Qiskit's QASM Simulator as opposed to on an actual quantum device. We worked mostly with simulations for a variety of reasons: 1), the state of the art physical devices have a small number of qubits, limited circuit depth, and are highly error prone; 2), we only have access to a few of IBM's publicly-available devices (see below), which often have long submission queues; and 3), our gradient descent method requires a high volume of circuit executions for each training data point, and the number of executions would quickly become unmanageable for a real device. In contrast, the QASM Simulator allows us to create large circuits with many executions without sacrificing accuracy of measurement, giving us more flexibility to explore the viability of QML in bioinformatics in ideal conditions. That said, we are interested in applying our QNN models in real conditions as well, so we decided to perform a subset of our testing on physical quantum computers.

##### *4d. IMPLEMENTATION ON REAL DEVICES*

In addition to the QASM Simulator, IBM offers cloud access to a number of real quantum devices for free. We have therefore supplemented our extensive testing on QASM Simulator with experimentation on IBM Quantum's open-access computers. All of the publicly-available computers are 5-qubit systems, circuits are limited to a quantum volume[1] of 32, and we have a maximum of 8,192 shots per simultaneous job submission. We've adapted our QNN models accordingly by subselecting four primary features from the dataset(s), reducing the number of layers in the QNN circuit, and paring our test dataset(s) down to a select few test cases to run sequentially.

For each modified QNN structure, we trained models on the appropriate feature groups using the simulator before classifying a limited number of test inputs on a real device. We split training and classification between simulated and actual devices, respectively, for a number of reasons. First, training (especially gradient descent) requires a large number of circuit executions in parallel, and that number far exceeds the 8,192 shots cap for job submissions to the public devices. Additionally, the number of public devices is quite limited, and all users of job submission must wait in the same global queue. Each data entry requires a different circuit construction because data must be encoded as rotation gates, so the hundreds of jobs needed for training would lead to impractical wait times. Finally, the error rates of the real device are much larger than those of the simulator, meaning the performance accuracy for a best model trained on a real device is expected to be lower than that of a model trained on the simulator. Training on simulators with low error rates guarantees a higher accuracy of the underlying trained model when the predictions are run on an actual quantum device.

Since our cQNN simulation code was written using Qiskit, changing from running on the QASM Simulator to one of IBM's publicly-available devices required only minor adjustments. We modified the custom-written class for the cQNN, adding a backend option to be able to switch between the QASM Simulator and cloud devices. We accessed the cloud devices via the IBM Quantum Lab virtual interface by uploading our cQNN code and running pre-trained models directly in the online notebook. We first loaded our IBM Quantum account into the notebook as the desired provider, then selected a backend from one of the 7 publicly-available systems, choosing the least busy device. Once we configured our modified cQNN class to use our selected backend, we executed our models on real devices. We ran each method with 1000 shots each. Depending on the length of the device access queue, using the cQNN to classify 9 test data points took anywhere from a couple of minutes to a couple of hours.

#### Margin Quantum Neural Network

The margin classifier was proposed by PennyLane in early 2020 and recommended by AWS. Detailed explanation as well as implementation can be found at https://pennylane.ai/qml/demos/tutorial_multiclass_classification.html . In brief, a quantum circuit is defined for each of the + and - classes. For each circuit, |F| number of features can be encoded into CEIL(log2|F|) qubits by amplitude encoding, a series of layer operations $L_l$ is applied to the qubits, a Pauli-Z measurement is made to a qubit, and lastly a bias term is added to yield the circuit output, $O_+$ or $O_-$. For the set of already-trained circuits, the binary classification

[1] "The largest random circuit of equal width and depth that the computer successfully implements," Measuring Quantum Volume (qiskit.org)

is made by the difference $O_+$ - $O_-$; for example, the positive label is predicted if the difference is positive. Each layer operation $L_l$ consists of single-qubit rotation ($\omega_{xq}$, $\omega_{yq}$, $\omega_{zq}$) for each qubit q, follow by two-qubit CNOT operations $CNOT_{q,mod(q+1, |q|)}$, between adjacent qubits. The rotation angles and bias terms are the hyperparameter of the circuit, and are trained by the minibatch with the cost computed from a loss function of ($O_{not\ l}$ - $O_l$ + margin) for a training point with label l. We note that ROC can be calculated by other (non-zero) thresholds of the difference ($O_+$ - $O_-$).

#### Weighted Margin Quantum Neural Network

Unlike the cQNN and qisQNN, we found that the aforementioned mQNN failed to converge under minibatch, Fig 5 (left) in the main text. We assert that such failure is due to the class imbalance of the difficult dataset. We also found by simple PCA and k-means that there are 3 clusters with different +/- class imbalance ratio. We perform 1:1 label-cluster-stratified split of the gold set data into training and validation portions. We performed a simple modification of the code from https://pennylane.ai/qml/demos/tutorial_multiclass_classification.html to execute without the minibatch and without unweighted cost. In particular, we train with all data points of the training set with $n_+$ positives and $n_-$ negatives, and re-weight the loss from each + or - training points with weights $w_+$ or $w_-$ such that $n_+w_+ = n_-w_-$. We found that the convergence is improved, Fig 5 (right) in the main text.

#### Molecular dynamics simulation

Molecular dynamics were performed with a pipeline featuring the high-performance GROMACS 2021.3 compiled in double precision (Berendsen et. al., 1995; Lindahl et. al., 2001; Van Der Spoel et. al. 2005; Hess et. al., 2008; Pronk et. al., 2013; Páll et al., 2015; Abraham et. al., 2015). Shorter 0.4ns simulations were performed for various ligands with low binding affinities estimated by AutoDock Vina (Trott et. al. 2010). Longer simulations (10ns) were performed on selected ligands, including the ligand named X77 that was co-crystallized with SARS-CoV-2 $3CL^{pro}$.

In preparation for GROMACS simulation, the receptor PDB files were processed using the “pdb4amber” function in Ambertools 2.0 (Morris et. al., 2009) to remove water molecules, add hydrogen bonds, and remove non-protein components. These processed PDB files provide initial receptor coordinates for GROMACS modeling with the AMBER03 force field (Duan et. al. 2003; Sorin et. al. 2005; Bremer et. al., 2020; Alvarado et. al., 2020). The starting ligand pose is prepared for GROMACS from the best docked pose estimated by AutoDock Vina and processed by Open Babel (O’Boyle et. al., 2011) and ACPYPE (Sousa da Silva et. al., 2012; Batista et. al., 2006; Wang et. al. 2004; Wang et. al. 20006) with General Amber Force Field (GAFF). The complex is solvated with the TIP3P model and neutralized by added ions.

### Supplementary Figures

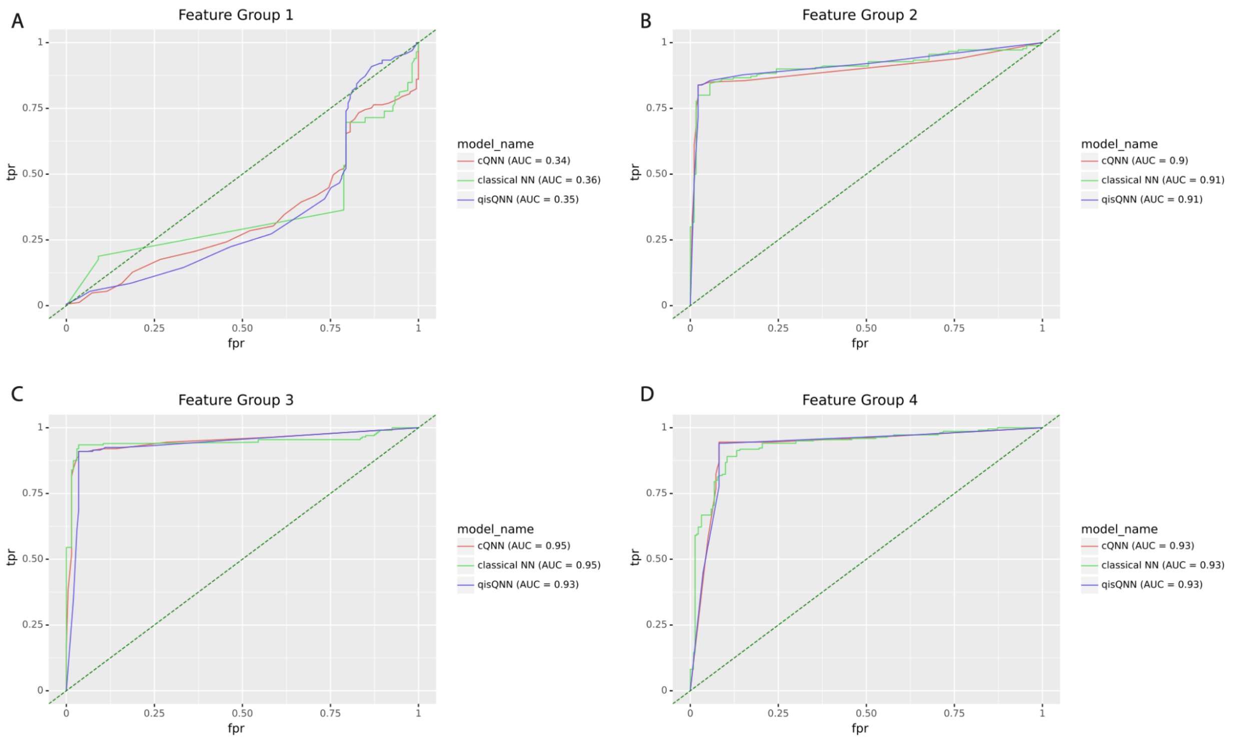


**Figure S1.** ROC curves (tpr = true positive rate vs. fpr = false positive rate) for the four feature groups of the GenoDock analysis applied to the leave-out GenoDock test dataset (a compilation of ExAC and TCGA datasets as described in the Supplementary Methods). The three curves in each panel indicate results for the cQNN, the qisQNN and a classical neural network (multilayer perceptron model implemented using Python's scikit-learn package) matched with the QNNs in terms of the number of parameters, and the corresponding areas under the curve (AUCs) are indicated in the legend. (A) Feature group 1: SNV features only (4 features in total); (B) Feature group 2: SNV + Structure features - Distance feature (7 features in total); (C) Feature group 3: SNV + Ligand features + Binding site "on/off" feature (10 features in total); (D) Feature group 4: SNV + Structure + Ligand features (13 features in total).

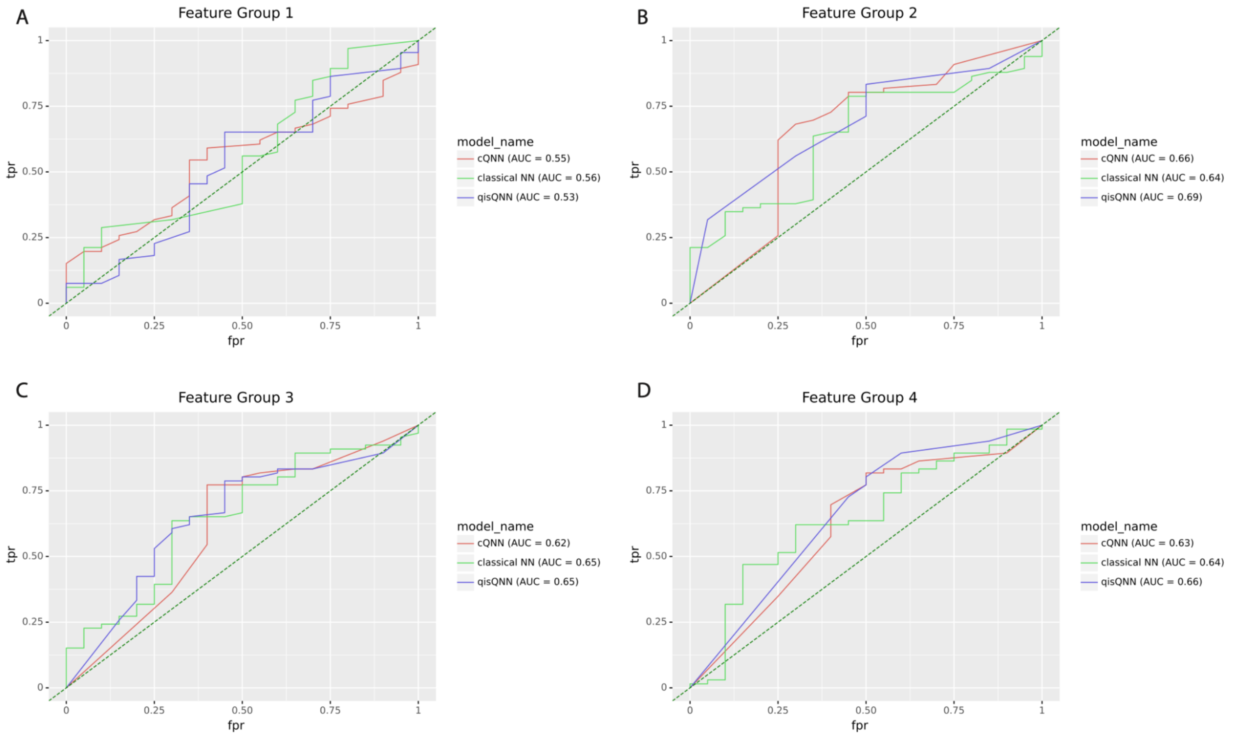


**Figure S2.** ROC curves (tpr = true positive rate vs. fpr = false positive rate) for the four feature groups of the GenoDock analysis applied to the Platinum dataset. The three curves in each panel indicate results for the cQNN, the qisQNN and a classical neural network (multilayer perceptron model implemented using Python's scikit-learn package) matched with the QNNs in terms of the number of parameters, and the corresponding areas under the curve (AUCs) are indicated in the legend. (A) Feature group 1: SNV features only (4 features in total); (B) Feature group 2: SNV + Structure features - Distance feature (7 features in total); (C) Feature group 3: SNV + Ligand features + Binding site "on/off" feature (10 features in total); (D) Feature group 4: SNV + Structure + Ligand features (13 features in total).

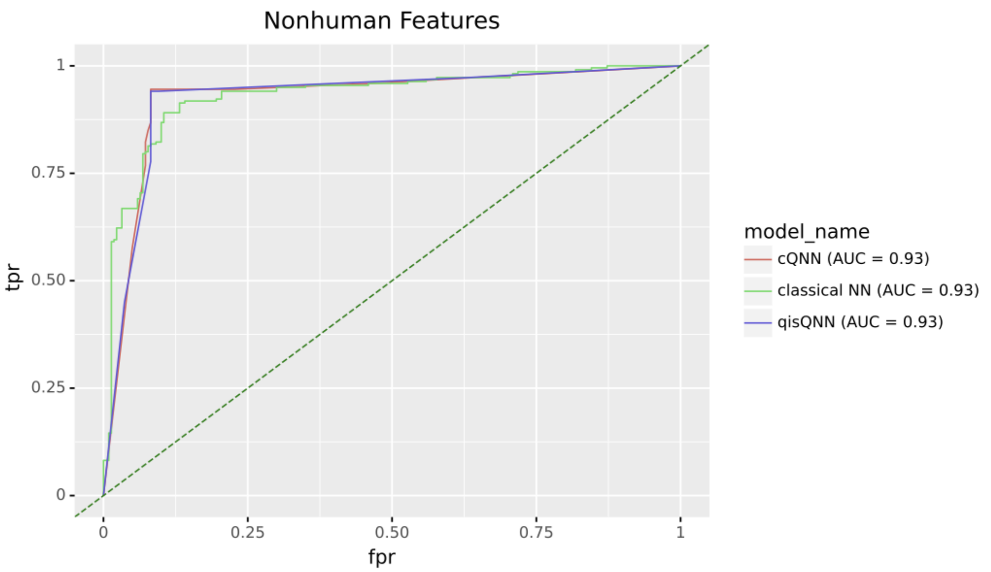


**Figure S3.** ROC curves (tpr = true positive rate vs. fpr = false positive rate) for the nonhuman (i.e. viral) feature group applied to the leave-out GenoDock test dataset (a compilation of ExAC and TCGA datasets as described in the Supplementary Methods). The three curves in each panel indicate results for the cQNN, the qisQNN and a classical neural network (multilayer perceptron model implemented using Python's scikit-learn package) matched with the QNNs in terms of the number of parameters, and the corresponding areas under the curve (AUCs) are indicated in the legend.

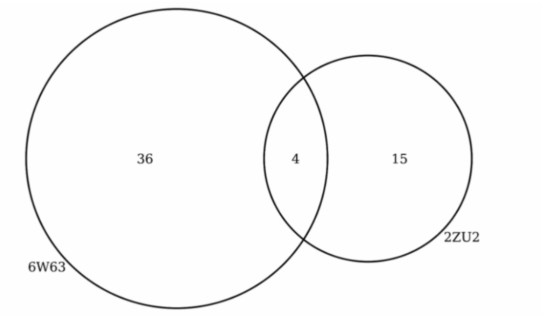


**Figure S4.** Comparison of AutoDock binding affinity. Venn diagram illustrating the number of ligands with AutoDock affinity ≤ -9.5 kcal/mol for the 6W63 and 2ZU2 receptors.

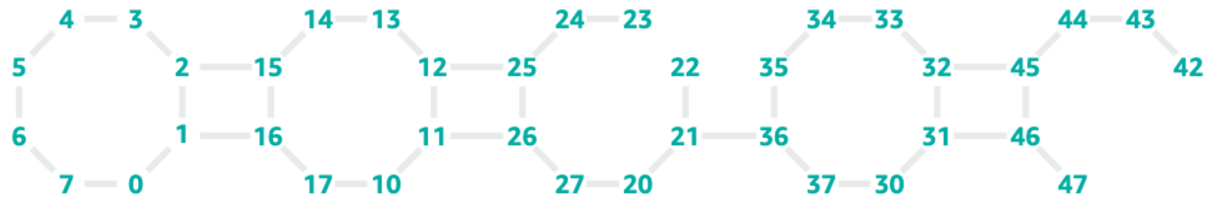


**Figure S5.** Qubit connectivity of the Rigetti system from Amazon Web Service.
